# TRIM24 directs replicative stress responses to maintain ALT telomeres via chromatin signaling

**DOI:** 10.1101/2024.10.18.618947

**Authors:** Daein Kim, Ragini Bhargava, Shih-Chun Wang, Doohyung Lee, Riya Patel, Sungtaek Oh, Ray W. Bowman, Chan Hyun Na, Roderick J. O’Sullivan, Kyle M. Miller

## Abstract

An inability to replicate the genome can cause replication stress and genome instability. Here, we develop BLOCK-ID, a proteomic method to identify and visualize proteins at stressed replication forks. This approach successfully identified novel mediators of the replication stress response, including the chromatin acetylation reader protein TRIM24. In validating TRIM24 function, we uncovered its crucial role in coordinating Alternative Lengthening of Telomeres (ALT), a cancer-specific telomere extension pathway involving replication stress. Our data reveal that TRIM24 is directed to telomeres via a p300/CBP-dependent acetylation chromatin signaling cascade, where it organizes ALT-associated PML bodies (APBs) to promote telomere DNA synthesis. Strikingly, we demonstrate that when artificially tethered at telomeres, TRIM24 can stimulate new telomere DNA synthesis in a SUMO-dependent manner, independently of p300/CBP or PML-dependent APBs. Thus, this study identifies a TRIM24 chromatin signaling pathway required for ALT telomere maintenance.

## Main

Faithful transmission of genetic and epigenetic material in eukaryotes requires the assembly of replicated DNA into chromatin, which compacts and organizes the genome through the formation of nucleosomes by histone proteins^1^. Histone proteins are highly modified by post-translational modifications (PTMs) that govern their cellular functions including during DNA replication. Replication stress is a major oncogenic driver that causes mutagenesis, chromosomal rearrangements, and genome instability^2,3^. Proteomic profiling of stressed replication forks has revealed an extensive network of DNA damage response (DDR) factors that respond to insults during DNA replication. These approaches include hydroxyurea (HU)-induced fork stalling by iPOND^4,5^ and TOP1 inhibitor-induced fork stress by Nascent Chromatin Capture (NCC) ^6^ have revealed a systemic reorganization of chromatin at stressed replication fork-associated. Other studies have also shown at stressed forks the assembly of heterochromatin^7^ and recruitment of chromatin remodeling factors to stabilize stressed forks and protect against transcription-replication conflicts^8^. Despite these advancements in proteomic methodologies that primarily rely on nucleotide analog integration into nascent DNAs^4,6,9^, an incomplete inventory of the stressed replication-specific responders, particularly epigenetic regulators, limits our understanding of mechanisms that respond to replication stress.

Here, we developed BLOCK-ID that combines the LacO/LacI replication barrier system with the proximity-dependent biotin identification method with proteomics and fluorescence microscopy to identify spatially associated DDR and chromatin factors at blocked replication forks. Using BLOCK-ID, we identified new proteins enriched at blocked replication forks, including the bromodomain (BRD) chromatin reader protein, Tripartite Motif containing 24, TRIM24. Our functional analyses uncovered an unexpected but crucial role for TRIM24 in mediating Alternative Lengthening of Telomeres (ALT), a pathological telomere extension mechanism detected in advanced cancer subtypes linked with mutations in chromatin modifiers, including ATRX-DAXX chromatin remodeling complex and histone H3.3^10,11^. The ensuing alterations in chromatin dynamics allow the build-up of pronounced replicative stress that stimulates HDR mechanisms like break-induced replication (BIR), which repairs and hyper-extends telomeres in those cancer cells^12–14^. These activities that extend telomeres have been visualized within unique liquid-liquid phase-separated (LLPS) nuclear compartments termed ALT-associated PML Bodies (APBs), which typically encompass 2∼5 telomeres^15^ and whose formation relies on multivalent contacts between SUMOylated proteins and SUMO-interaction motifs (SIM) present within PML and mediators of ALT-HDR. APBs are enriched with DNA damage response (DDR) factors, including the MRN complex, BRCA1, RAD51, RAD52, and the BTR (BLM, TOP3A and RMI1/2) complex ^16–18^. This composition provides an optimal environment for telomere extension by sequestering telomeres and DDR factors. Indeed, nascent telomere synthesis is predominantly observed within APBs^19^. Consistent with the purported essential role of APBs in ALT, disruption of their core constituent PML attenuates ALT-HDR, leading to gradual telomere attrition^20^.

This study demonstrates that TRIM24 fulfills crucial roles in maintaining telomeres in ALT cancer cells. TRIM24 deficient cells exhibit features consistent with impaired ALT, most notably impaired APB formation, robust telomere shortening and diminished integrity due to elevated replicative stress. By screening known human histone acetyltransferases, we determined that TRIM24 is recruited to telomeres via CBP/p300 histone acetyltransferases that mediate the acetylation of histone H3 lysine 23. TRIM24 recognizes this acetylation signal at damaged telomeres through its bromodomain and once localized, stimulates SUMOylation on two key residues within TRIM24 by PIAS1, which promotes ALT activities at telomeres. Our analysis determined that even though these interactions are required for functional APB formation, we uncovered that forced tethering of TRIM24 to telomeres bypassed the requirement for p300/CBP and the key APB structural component PML for initiating new telomere synthesis. Thus, this study reveals a PTM-driven epigenetic signaling pathway involving histone acetylation recognition and SUMOylation of TRIM24 that establishes ALT-mediated telomere maintenance in cancer cells.

## Results

### BLOCK-ID identifies bromodomain proteins at obstructed replication forks

We sought to develop a system for detecting proteins in living cells that accumulate at obstructed replication forks without needing DNA synthesis-dependent labeling^4,6,9^. In human osteosarcoma cell line U2OS genomically harboring a 256 tandem repeat array of the lac operator (LacO) sequence (U2OS 256x), replication fork progression can be blocked by expressing the lac repressor LacI as it tightly binds to the integrated LacO array^21,22^. We introduced the promiscuous biotin ligase BirA*^23^ fused to LacI (BirA*-LacI) into U2OS 256x cells to enable biotin labeling of proteins associated specifically with blocked replication forks (Fig. 1a). We name this technique BLOCK-ID for Biotinylation of LacO array repliCation stress protein networ<u>K</u> <u>ID</u>entification. To avoid persistent replication fork blocking BirA*-LacI expression was controlled by doxycycline. Induction of BirA*-LacI and supplementation of media with exogenous biotin led to protein biotinylation, both in bulk measurements by Western blotting and within foci where BirA*-LacI localized to the LacO array (BLOCK-ID foci) (Fig. 1b). By immunofluorescence (IF), we observed the accumulation of phospho-ATR kinase (pATR -S428), BRCA1, RPA2, and γH2AX confirming the activation of replication stress responses at blocked replication forks at BLOCK-ID foci (Fig. 1c; see methods).

**Fig. 1:**
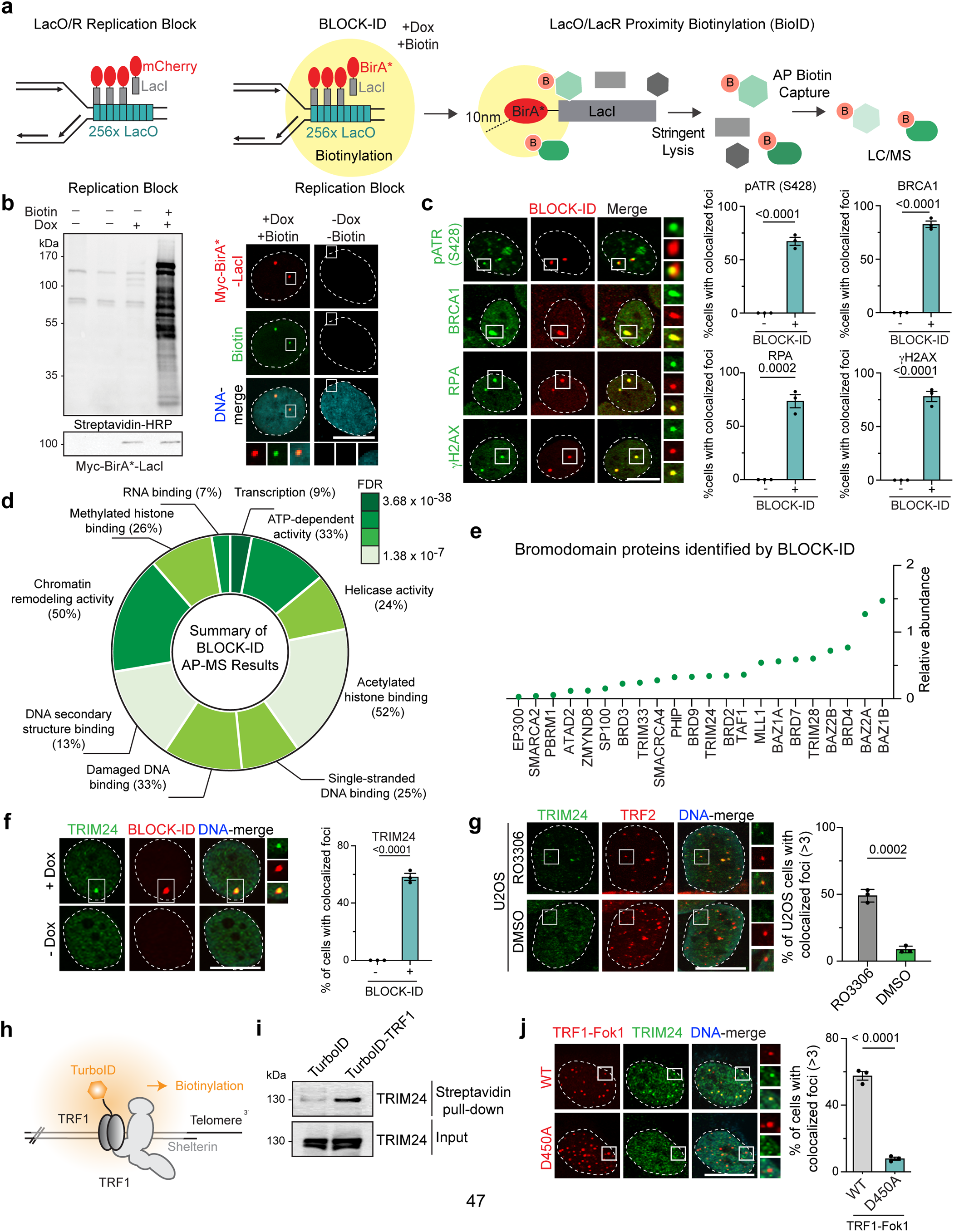
Stressed replication fork profiling by BLOCK-ID identifies TRIM24. **a,** Schematic of BLOCK-ID. The yellow circle indicates the biotinylation range of BirA* (10 nm). Biotinylated proteins are isolated by biotin affinity pulldown (AP) and analyzed by liquid chromatography mass spectrometry (LC/MS). **b,** Western blot of proteins biotinylated by BLOCK-ID. Cells were treated with doxycycline (Dox) and/or biotin (50 μM) for 14 hrs. **(b, right)** representative immunofluorescence (IF) images by confocal microscopy showing the localization of biotinylated proteins at BLOCK-ID foci. **c,** Localization of replication stress markers at BLOCK-ID foci was analyzed using IF and confocal microscopy. Quantification is on the right, n=3, >100. **d,** Gene Ontology (GO) analysis of BLOCK-ID identified proteins. **e,** Plot depicting the relative abundance of bromodomain (BRD) proteins identified by BLOCK-ID. **f,** Localization of endogenous TRIM24 at BLOCK-ID foci was analyzed using IF and confocal microscopy. Quantification is on the right, n=3, >100. **g,** Localization of endogenous TRIM24 at telomeres was analyzed using IF and confocal microscopy. Telomeres were detected using TRF2 antibodies. Cells were treated with 9 μM of RO3306 (G2 arrest) or DMSO for 16 hrs. Quantification is on the right, n=3, >100. **h,** Telomere-specific biotinylation by TurboID-TRF1 in U2OS cells. The yellow circle indicates biotinylation range of TurboID. **i,** Western blot of lysates following streptavidin pull-down from U2OS cells expressing TurboID-TRF1 detects TRIM24. **j,** Localization of endogenous TRIM24 at ALT telomers indicated by TRF1-FokI WT or D450A foci in U2OS cells. Quantification is on the right, n=3, >100. Unless otherwise indicated, statistical analyses were performed with unpaired two-tailed T-test, error bars in graphs represent mean ± s.e.m., with P values indicated in the graphs. Dots on the graphs indicates independent experiments. N represents number of biologically independent experiments and indicated number represents total number of cells or treats analyzed per each experimental group. All scale bars in IF images, 10 μm.

Next, we performed a large-scale BLOCK-ID mass spectrometry-based screen. Here, experiments in growth-arrested cells and cells expressing BirA* alone were included to control for non-specific biotinylation locally at LacO/I repeats and globally throughout the cell. Our analysis identified a total of 591 proteins (Extended Data Table 1 and methods), including components of the DNA replisome and fork protection complexes (Extended Data Fig. 1a) and various DNA repair pathways (Extended Data Fig. 1b-c) including several previously identified by proteomics of HU-arrested replication forks^4,6^ (Extended data Fig. 1d-f). This analysis supports the utility of BLOCK-ID as a new method for identifying proteins associated with blocked replication forks. Through Gene Ontology (GO) analysis, DNA and chromatin proteins were also identified (Fig. 1d). Of those, proteins containing the acetylation reader BRD were highly enriched, constituting 55% of all ubiquitously expressed human BRD-containing proteins (22 out of 40; Fig. 1e). BRD proteins act as “epigenetic readers” for acetylated lysines within both histone and non-histone proteins^24^ and play essential roles in DNA repair pathways^25–30^. However, their involvement in replication stress responses remain poorly understood^31^.

To address this, we assessed the recruitment of BRD proteins to blocked forks at BLOCK-ID foci by IF, where 12 out of the 22 tested BRD proteins displayed clear recruitment (Extended Data Fig. 2a; quantified in b). Furthermore, depletion of 10 out of the 22 BRD proteins led to >30% cell death upon HU treatment (Extended Data Fig. 2c, the efficiency of siRNA depletion is provided in d). Comparing results from these two different assays identified BAZ1B, BRD9, TRIM24 and ZMYND8 as regulators of the replication stress response and showcased the value of BLOCK-ID in identifying proteins associated with blocked replication forks, including chromatin regulatory factors.

### Mobilization and function of TRIM24 at blocked replication forks

TRIM24 was of particular interest given that its poorly characterized in DNA damage responses, including during DNA replication, and expression levels correlate with the progression of various cancers^32,33^. TRIM24 is a member of a large TRIM protein family characterized by the conserved N-terminus sequentially comprised of a RING finger domain, B-box type zinc finger domains, and a coiled-coil (CC) domain^34^. TRIM24 features a tandem plant homeodomain (PHD) and BRD reader domain in the C-terminus. TRIM24 has been characterized as a transcription coregulator^35^ and a ubiquitin E3 ligase for p53^36^, as well as a DNA double-strand break repair-associated factor from our previous screens^37,38^

Using a validated antibody against endogenous TRIM24, we observed its localization at BLOCK-ID foci (Fig. 1f; antibody specificity validated in Extended Data Fig. 3a-b). We also performed SIRF (in situ protein interactions at nascent and stalled replication forks) ^30,39^ with BLM as a positive control that detected an association of TRIM24 with stalled replication forks in HU-treated conditions (Extended Data Fig. 3c). Recruitment of TRIM24 to blocked forks required the replication stress kinase ATR but was independent of other DNA damage associated signaling factors including the kinase ATM or Poly(ADP) ribose polymerase PARP1, consistent with recruitment of TRIM24 to BLOCK-ID foci being a response to replication stress (Extended Data Fig. 3d) ^40^. Depletion of TRIM24 by siRNA elevated pATR under normal growth conditions compared to siControl (siCTRL) cells (Extended Data Fig. 3e). When siCTRL and siTRIM24 cells were treated with the replication stress-inducing agent aphidicolin, replication stress signals (i.e. pATR and γH2AX) were maintained in cells deficient of TRIM24 (Extended Data Fig. 3e). All together, these data support TRIM24 as a replication stress response factor.

### Specific association of TRIM24 with telomeres in ALT cancer cells

We noticed that endogenous labeling of TRIM24 in cycling U2OS cells displayed a punctate staining pattern (Fig. 1f). Given that U2OS cells are a well-characterized cancer cell line that relies on the ALT mechanism for telomere extension, which represents a naturally occurring replication-stress pathway, we hypothesized that TRIM24 might localize to telomeres in ALT cancer cells. Indeed, a previous report described TRIM24 interacting with PML^41^, a key component of ALT-associated PML bodies (APBs), where telomere extension occurs during G2-M cell cycle phase^13^. Thus, we analyzed telomere recruitment of TRIM24 in G2-arrested U2OS cells using the CDKi, RO3306. Indeed, endogenous TRIM24 localized more efficiently to telomeres (Fig. 1g). TRIM24 was also robustly detected following western analysis of proteins captured by proximity-dependent biotinylation with TRF1-TurboID (Fig 1h-i). Neither endogenous TRIM24 nor overexpressed GFP-TRIM24 were observed at telomeres in asynchronous or G2-arrested telomerase-positive HeLa cells with long telomeres (HeLa LT) (Extended Data Fig. 4a-b). Replicative stress and DNA breaks stimulate the ALT-associated homology-directed repair mechanism. Thus it was notable that TRIM24 localization to telomeres was enhanced by pyridostatin (PDS) treatment (Extended Data Fig. 4c), a G4-quadruplex stabilizing ligand known to perturb telomere replication^42^. Consistently, the enrichment of TRIM24 at telomeres was observed following the generation of telomere-specific DNA breaks with TRF1-FokI in U2OS cells (Fig 1j), but not HeLa LT cells (Extended Data Fig. 4b). The apparent selective and stimulus-responsive localization of TRIM24 with telomeres led us to conclude that TRIM24 could represent a mediator of ALT.

### TRIM24 is required for productive ALT

We evaluated the impact of TRIM24 knockdown on distinctive aspects of ALT telomere. By combining immunofluorescence (IF) for PML with telomeres, we observed that TRIM24 depletion reduced the frequency of ALT-associated PML bodies in U2OS cells (Fig. 2a). Similarly, using the native FISH-based ssTelo assay that detects C-rich telomeric ECTRs^20^, including partially single-stranded C-circles, TRIM24 depletion reduced the number of ssTelo signals in U2OS cells (Fig. 2b). Notably, the reduction in ssTelo observed after TRIM24 depletion was comparable to removing BLM ^17,20,43,44^, an essential regulator of ALT which suppresses excessive telomere recombination. By performing Chromosome-Orientation FISH (CO-FISH) assay on metaphase chromosomes, we found that whereas BLM deficiency elicits high rates of telomere recombination and telomere-sister chromatid exchanges (t-SCEs) (Fig. 2c, indicated with carets), TRIM24 depletion decreased t-SCE frequency (Fig. 2c). The increased t-SCE frequency in BLM-depleted cells was suppressed to normal levels after co-depletion of TRIM24 (Fig. 2c). Therefore, these data indicate that TRIM24 may contribute to several aspects of telomeres (i.e. APB formation, telomere recombination, etc.) promoting recombination that are essential for productive ALT and maintaining telomere length in those cancer cells.

**Fig. 2:**
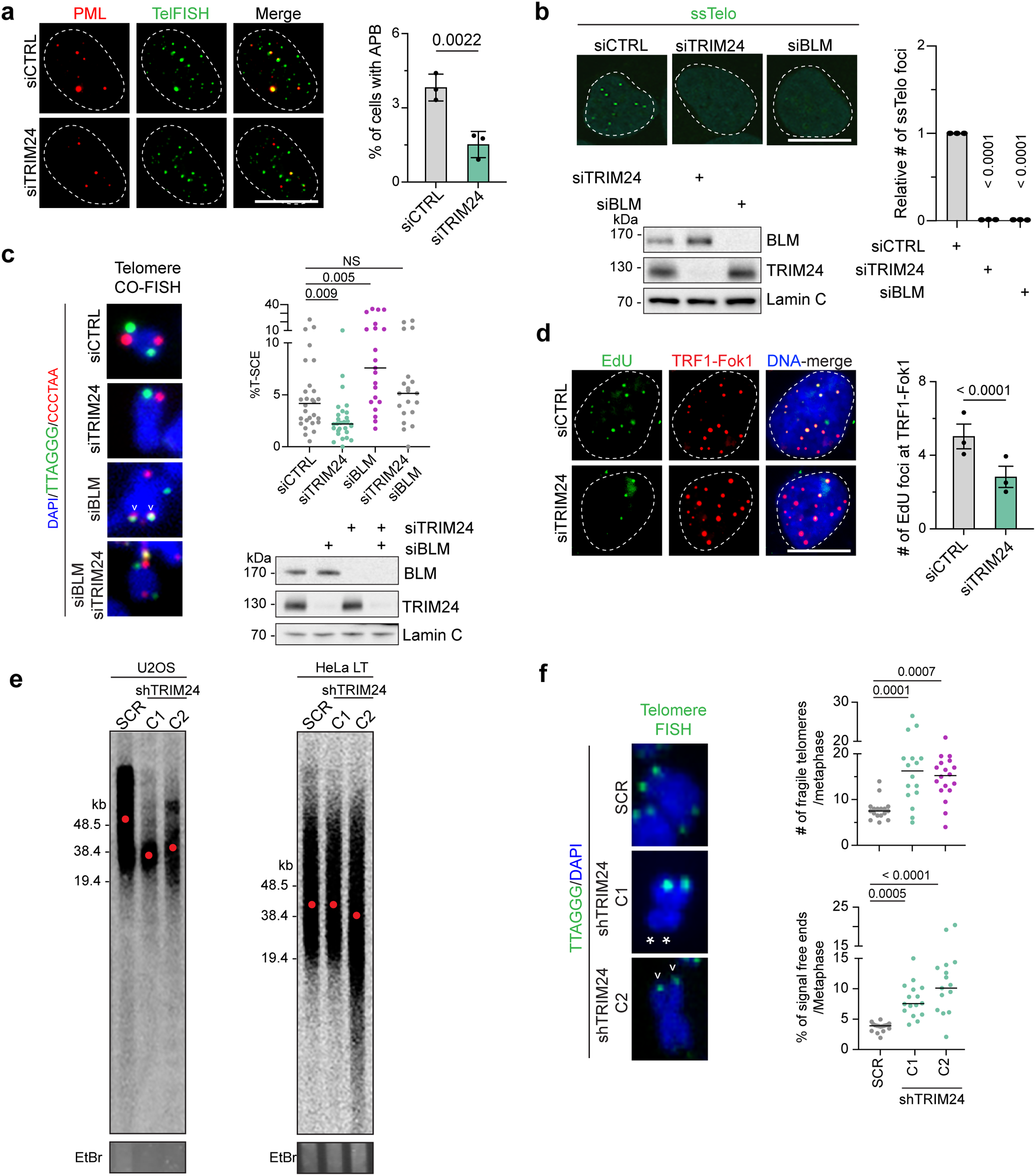
TRIM24 promotes ALT telomere maintenance. **a,** APBs were analyzed in U2OS cells treated with CTRL or TRIM24 siRNAs. Telomeres were detected using TRF2 antibodies and imaged with fluorescence microscopy. Quantification on the right, n=3, > 2500. **b,** Detection of ssTelo foci by FISH in U2OS cells following treatment with siRNAs for CTRL, TRIM24 or BLM. Western blot analysis shows depletion of the indicated proteins using siRNAs. Lamin C is the loading control. Quantification with one-way ANOVA is on the right, n=3, > 450. **c,** Analysis of CO-FISH in U2OS cells following TRIM24, and/or BLM depletion. Carets indicate T-SCEs. Quantification with one-way ANOVA is on the top right, n=3, > 30 metaphase spreads. Validation of BLM, and TRIM24 depletion using the indicated siRNAs by Western blotting of lysates from these treatments is shown on the bottom right. **d,** Nascent telomere synthesis detection by EDU incorporation was analyzed in U2OS cells following TRIM24 depletion. TRF1-FokI indicates telomeres. Quantification is shown on the right, n=3, >150. **e,** Telomere length analysis by PFGE. TRIM24-depleted U2OS and HeLa LT cells were generated by stable expression of two independent shRNAs (C1 and C2) targeting the coding region of TRIM24. SCR is non-targeting shRNA. Ethidium bromide (EtBr) is the loading control. Red dots indicate the mean telomere length. **f,** Analysis of fragile telomeres and signal free ends was performed using telomere FISH on metaphase spreads. Cells from **e** were analyzed. Carets indicates fragile telomeres and asterisks indicate signal free ends. Quantification with one-way ANOVA is provided on the right, n=2, >30 metaphase spreads. Unless otherwise indicated, statistical analyses were performed with unpaired two-tailed T-test, error bars in graphs represent mean ± s.e.m., with P values indicated in the graphs. NS indicates no statistical significance. Dots on the graphs indicates independent experiments. N represents number of biologically independent experiments and indicated number represents total number of cells, or treats analyzed per each experimental group. All scale bars in IF images, 10 μm.

To determine the impact of TRIM24 deficiency on ALT-mediated telomere elongation, we first turned to the TRF1-FokI system, which enables the synchronous and robust induction of nascent telomere DNA synthesis^13,14^. Here, nascent telomere DNA synthesis can be quantified by monitoring the accumulation of the nucleotide analog, Ethynyl-2’-deoxyuridine (EdU), at TRF1-FokI generated DNA breaks^12^. Using this assay, ∼50% fewer telomeres displayed focal accumulation of EdU within 72hrs of TRIM24 loss (Fig. 2d). To evaluate the impact of TRIM24 loss on telomere length directly, TRIM24 was stably depleted in U2OS and HeLa LT cell lines using short hairpin RNAs (shRNA) (Extended Data Fig. 5a). We confirmed by flow cytometry that TRIM24 loss does not appreciably alter the cell cycle in these cell lines (Extended Data Fig. 5b) even though the ALT+ cells displayed elevated phosphorylation of the replication stress marker serine 33 on RPA2 (pRPA2-S33) (Extended Data Fig. 5a). Following the sustained culture of WT and TRIM24-deficient U2OS and HeLa LT cells, telomere length was directly measured by pulsed-field gel electrophoresis of telomere restriction DNA fragments followed by Southern blotting with telomere probes. This analysis revealed substantial telomere shortening in TRIM24-deficient U2OS cells but not TRIM24-deficient HeLa LT cells (Fig. 2e). Telomere FISH analysis on metaphase chromosome spreads prepared from control and TRIM24-deficient U2OS cells. revealed a greater frequency of telomere fragility in TRIM24 deficient cells (Fig. 2f, indicated with carets), a phenotype associated with incomplete replication and/or replicative stress at telomeres. Moreover, the incidence of chromosome ends without FISH signals (Signal free ends (SFEs)) was markedly increased (Fig. 2f, indicated with asterisks), indicating substantial telomere loss in TRIM24 deficient U2OS cells, which is consistent with our telomere length measurements of bulk cultures. Notably, the specific impact of TRIM24 on telomere length and ssDNA at telomeres in ALT cancer cells was confirmed by the same analyses of telomeres from matched isogenic LM216J (ALT+) and LM216T (ALT-) cell lines. TRIM24 depletion reduced telomeric ssDNA in LM216J (ALT+) cells and telomere length analysis also demonstrated a rapid decrease in telomere length in LM216J (ALT+) but not LM216T (ALT-) cell lines (Extended Data Fig. 5c-f). TRIM24 deficient LM216J cells also exhibited elevated pRPA33, as well as impaired APB formation and nascent telomere DNA synthesis (Extended Data Fig. 5c,g-h). These data unequivocally demonstrate that TRIM24 is required for ALT-mediated telomere length maintenance.

### CBP and p300 recruit TRIM24 to telomeres to facilitate ALT

We next sought to determine the molecular requirements for TRIM24 recruitment to ALT telomeres. Using a series of domain deletion derivatives of TRIM24, localization of TRIM24 to telomeres was determined to be dependent on the PHD and BRD domains (Fig. 3a). The PHD and BRD of TRIM24 are single “reader” motifs, which have been shown to bind a chromatin signature of unmethylated H3K4 and histone H3 acetylated at lysine 23 (H3K23ac) ^33^ involving gene regulation in breast cancer cells^33,45^. Since TRIM24 recruitment to damaged telomeres depended on the PHD and BRD domains, we reasoned that histone modifications may mediate the interaction between chromatin and TRIM24 at telomeres in ALT cancer cells. To test this possibility, we examined whether any of 12 major histone acetyltransferases (HATs) might promote TRIM24 recruitment to telomeres. We depleted each of these HATs in U2OS cells and monitored the localization of TRIM24 to TRF1-FokI telomeric breaks. Notably, depleting CBP and p300 markedly reduced telomere localization of TRIM24 (Fig. 3b; Screen shown in Extended Data Fig. 6a-c). TRIM24 telomere localization was ablated following treatment of cells with the CBP and p300 HAT activity inhibitor A-485^46^ (Extended Data Fig. 7a). Importantly, depletion of either CBP or p300 did not noticeably alter the mRNA or protein level of TRIM24 (Extended Data Fig. 7b), suggesting a direct role for the HAT activity of CBP and p300 in directing TRIM24 to telomeres in ALT cancer cells. Indeed, CBP and p300 localized to TRF1-FokI induced telomere DSBs in U2OS cells (Fig. 3c), but not in non-ALT, telomerase-positive HeLa LT cells (Extended Data Fig. 7c).

**Fig. 3:**
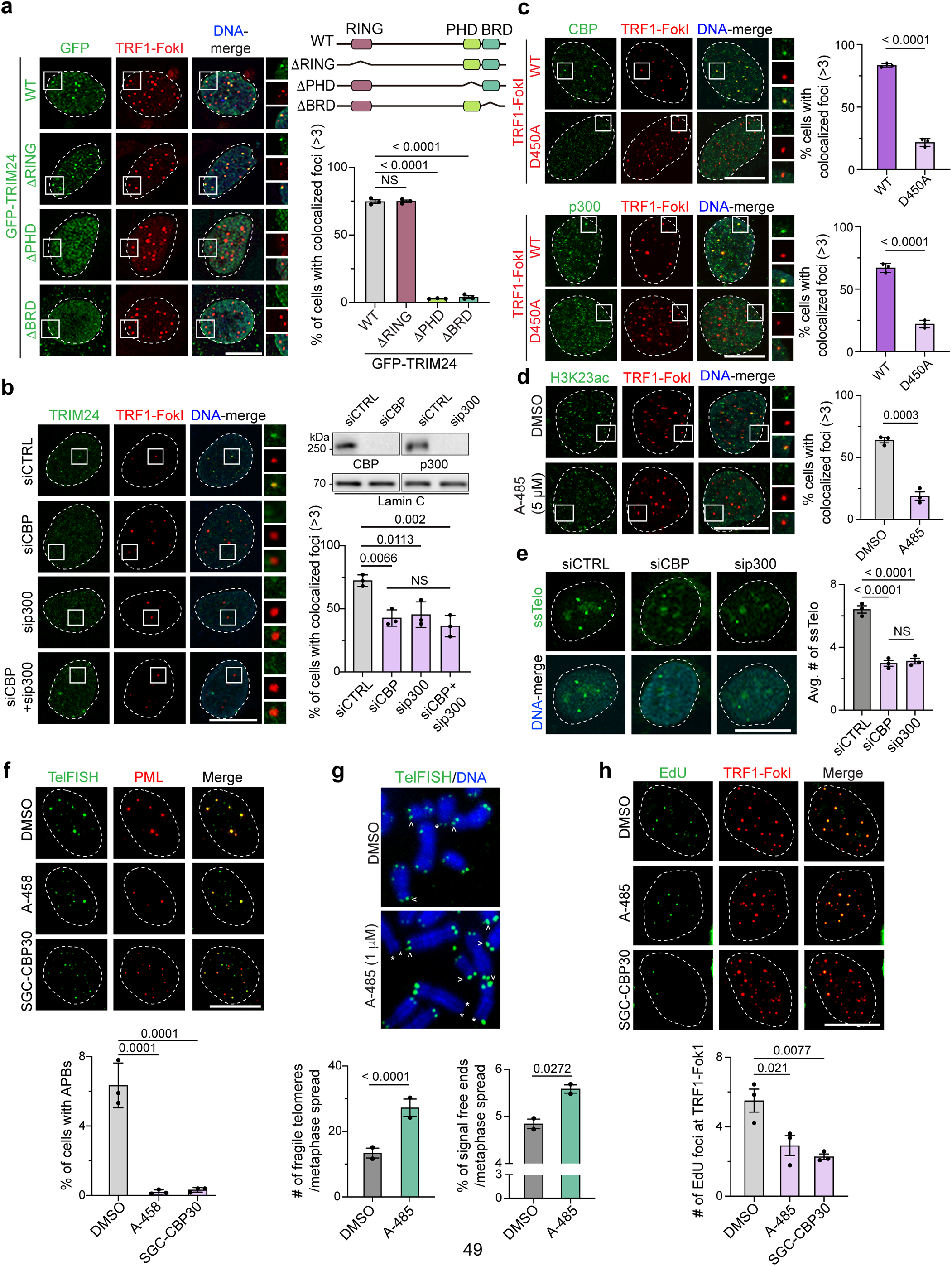
CBP and p300 participate in ALT by directing TRIM24 to telomeres. **a,** Colocalization of GFP tagged TRIM24 WT and deletion variants with TRF1-FokI foci in U2OS cells was analyzed by confocal microscopy. Scheme of WT TRIM24 domains and deletion variants is shown on the top right. Quantification is on bottom right, n=3, > 100. **b,** Experiments in **a** were performed in U2OS cells following treatment with siRNAs targeting CTRL, CBP and/or p300. Western blot of lysates from treatments analyzed with indicated antibodies is on the top right. Quantification of **b** is on the bottom right, n=3, > 96. **c,** Colocalization of endogenous CBP and p300 with TRF1-FokI WT and D450A foci in U2OS cells was analyzed using IF and confocal microscopy. Quantification with unpaired two-tailed T-test is shown on the right, n=3, >105. **d,** Telomere localization of H3K23ac following TRF1-FokI induction in U2OS cells was analyzed using IF and confocal microscopy. Cells were treated with 5 μM of A-485 or DMSO for 16 hrs prior to analysis. Quantification with unpaired two-tailed T-test is on the right, n=3, >400. **e,** Detection of ssTelo foci following depletion of CBP or p300 in U2OS cells was performed using confocal microscopy. Quantification is on the right, n=3 >420. **f,** APBs in U2OS cells treated with A-485 or SGC-CBP30 was analyzed. Cells were treated with 1 μM of A-485 or 1 μM of SGC-CBP30 for 3 days. Telomeres were detected using telomere FISH (TelFISH). Quantification is provided below, n=3, >2500. **g,** Analysis of fragile telomeres and signal free telomeres in U2OS cells treated with A-485 or DMSO. Cells were treated with 1 μM of A-485 or DMSO for 3 days. Carets indicate fragile telomeres and asterisks indicates signal free ends. Quantifications with two-tailed T-Test are provided below, n=2, >30 metaphase spreads. **h,** Nascent telomere synthesis was analyzed as in Fig. 2d for cells treated as in **f**. Quantification is provided below, n=3, > 150. Unless otherwise indicated, statistical analyses were performed with one-way ANOVA, error bars in graphs represent mean ± s.e.m., with P values indicated in the graphs. NS indicates no statistical significance. Dots on the graphs indicates independent experiments. N represents number of biologically independent experiments and indicated number represents total number of cells, or treats analyzed per each experimental group. All scale bars in IF images, 10 μm.

TRIM24 interactions with H3K23ac and H3K14ac have been reported^33^. Thus, we asked whether H3K23ac and/or H3K14ac are present at telomeres using U2OS cells expressing TurboID-TRF1. Using specific antibodies to these histone marks, both H3K23ac and H3K14ac were detected in the biotinylated protein fraction recovered from streptavidin pulldown in ALT cells. Notably, depleting CBP markedly diminished the level of H3K23ac level in the biotinylated protein fraction, whereas H3K14ac remained unchanged (Extended Data Fig.7d-e). In agreement with these biochemical data, we observed a substantial accumulation of H3K23ac following the induction of telomere DSBs with WT-TRF1-FokI (Extended Data Fig. 7f), which was diminished after treatment with A-485 (Fig. 3d). Previous studies reported that the MOZ/MORF protein complex catalyzes acetylation of H3K23 that is involved in gene regulation^47,48^. Following the depletion of BRPF1, a component of the MOZ/MORF complex, we observed reduced global levels of chromatin-associated H3K23ac and TRIM24 as expected (Extended Data Fig. 7g). However, depletion of either the HAT MORF or BRPF1 did not affect TRIM24 localization to telomeres, (Extended Data Fig. 6a-c and 7h-i). These data suggest the presence of a telomere-specific H3K23ac signal mediated by CBP and p300 that promotes TRIM24 localization to telomeres.

Considering the indispensable role of TRIM24 in ALT that we identified, we speculated that inhibition of the HAT activity of CBP and p300 might replicate the telomere phenotypes observed in TRIM24-deficient cells. Indeed, depleting CBP or p300 significantly decreased ssTelo signals in U2OS cells (Fig. 3e). Treatment with CBP/p300 inhibitors resulted in a pronounced increase in telomere fragility (indicated with carets) and telomere loss (indicated with asterisks) with a concomitant reduction in APB formation and telomere DNA synthesis (Fig. 3f-h). Collectively, these results have uncovered a chromatin signaling axis involving CBP/p300-TRIM24 that maintains telomere length regulation and telomere integrity in ALT cancer cells.

### Constitutive tethering of TRIM24 to telomeres promotes APB formation and enhances ALT

Given that TRIM24 telomere localization promotes ALT, we queried the effects of constitutive tethering of TRIM24 to telomeres. To do this, we fused the DNA-binding domain of the *S. pombe* protein Teb1 that recognizes TTAGGG human telomeric sequences^20,49^ to the C-terminus of TRIM24 (TRIM24-TebDB) and observed a complete colocalization of TRIM24-TebDB with the telomere binding protein TRF2 (Fig. 4a). Notably, constitutive tethering of TRIM24 via TebDB facilitated ALT activities, as indicated by increased ssTelo signals compared to control WT U2OS cells (Fig. 4b). Importantly, while CBP/p300 deficiency abolished ssDNA at telomeres consistent with its requirement for ALT, forced tethering of TRIM24 to telomeres in either CBP-depleted cells or A-485 treated cells rescued telomeric ssDNA formation (Extended Data Fig. 8a and Fig. 4b). These data indicate that even though CBP/p300 HAT activity is pivotal for initially recruiting TRIM24 to telomeres, it is dispensable for downstream events in ALT.

**Fig. 4:**
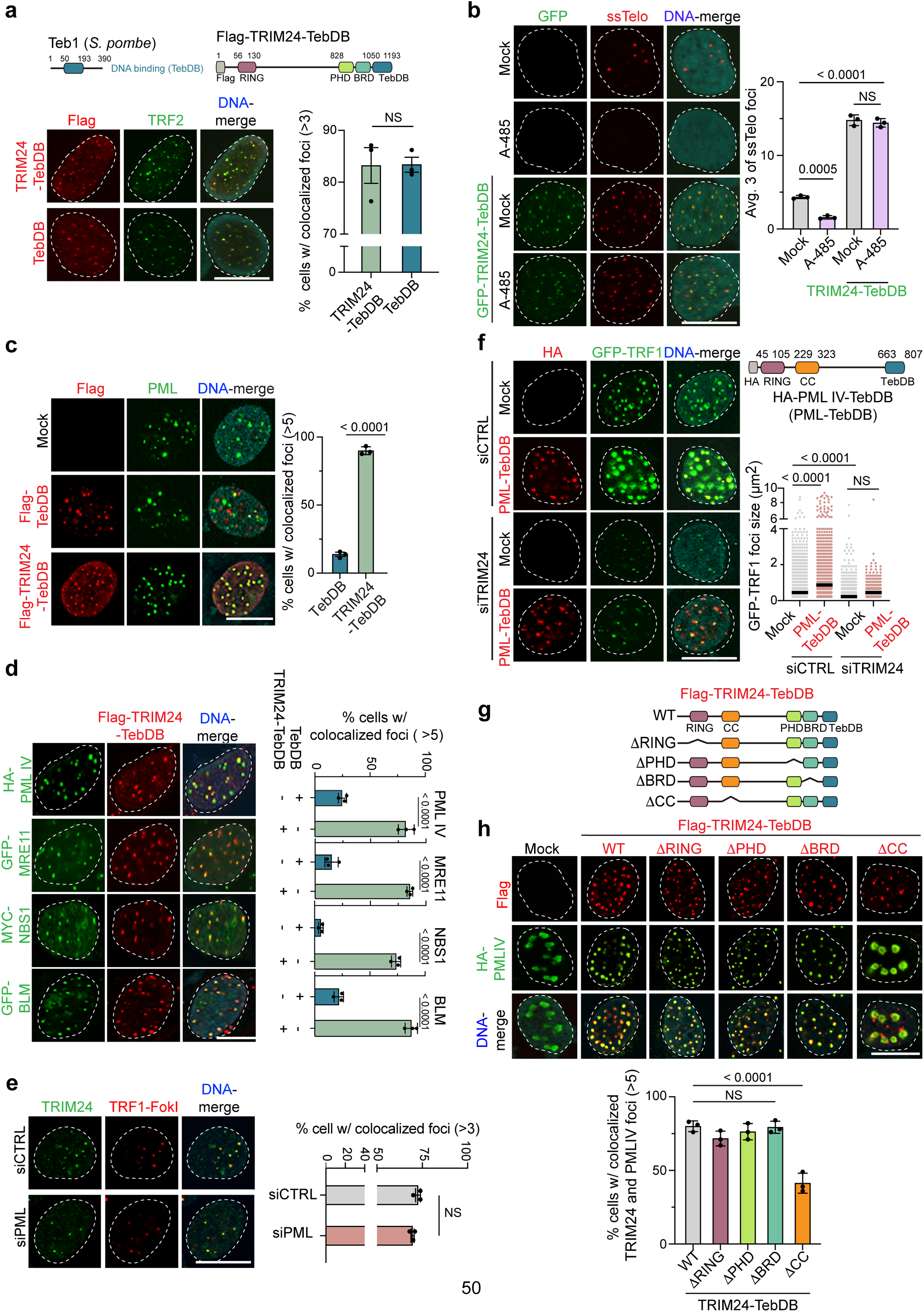
TRIM24 promotes APB formation at ALT telomeres. **a,** Localization of Flag-TebDB and Flag-TRIM24-TebDB to telomeres in U2OS cells was analyzed by IF and confocal microscopy. Illustration of Teb1 and TRIM24-TebDB is on the top. DNA binding domain of Teb1 (TebDB) is indicated in blue. TebDB was fused to the C-terminus of Flag-TRIM24 to generate Flag-TRIM24-TebDB. TRF2 labels telomeres. Quantification is provided on the right, n=3, >75. **b,** ssTelo foci were analyzed by FISH, IF and confocal microscopy in U2OS cells either treated or not-treated with 5 μM of A-485 for 72 hrs with or without expressing GFP-TRIM24-TebDB. Quantification with one-way ANOVA is provided on the right, n=3, >100. **c,** Colocalization of endogenous PML and Flag-TebDB or Flag-TRIM24-TebDB in U2OS cells was analyzed by IF and confocal microscopy. Quantification is shown on the right, n=3, >450. **d,** Colocalization of HA-PML IV, GFP-MRE11, Myc-NBS1, and GFP-BLM with Flag-TRIM24-TebDB was analyzed by IF and confocal microscopy. Quantification is on the right, n=3, >100. **e,** Localization of endogenous TRIM24 to telomeres in U2OS cells was analyzed following treatment with siCTRL or siPML. TRF1-FokI indicates telomeres. Quantification is provided on the right, n=3, >125. **f,** Telomere size in either mock-treated or HA-PML IV-TebDB (PML-TebDB) expressing U2OS cells following TRIM24 depletion was analyzed by IF and confocal microscopy. GFP-TRF1 marks telomeres. Schematic of HA-PML IV fused to TebDB (PML-TebDB) is shown on the top right. Quantification with one-way ANOVA is on the bottom right, n=3, >60. **g,** Domain structure of WT TRIM24 fused to TebDB and deletion variants. **h,** Colocalization of HA-PML IV with TRIM24 derivatives from **g** were analyzed by IF and confocal microscopy. Quantification with one-way ANOVA is provided below, n=3, >110. Unless otherwise indicated, statistical analyses were performed with unpaired two-tailed T-test, error bars in graphs represent mean ± s.e.m., with P values indicated in the graphs. NS indicates no statistical significance. Dots on the graphs indicates independent experiments. N represents number of biologically independent experiments and indicated number represents total number of cells, or treats analyzed per each experimental group. All scale bars in IF images, 10 μm.

This system provides a powerful experimental tool to dissect the sequence of molecular events dictated by p300-CBP-TRIM24 in ALT. Given the interaction between TRIM24 and PML and the role of PML in forming APBs, which are required for ALT^20,43^, we tested the recruitment of PML by TRIM24-TebDB to ALT telomeres. While TebDB displayed robust telomere localization, including at a few PML foci, TRIM24-TebDB telomere tethering resulted in a near complete localization of endogenous PML to TRIM24-TebDB bound telomeres compared to TebDB alone (Fig. 4c). Previous studies identified that the fourth splicing isoform of PML (PML IV) is essential for ALT^20,43^. Thus, we transfected HA-tagged PML IV into TRIM24-TebDB expressing cells to test its recruitment to telomeres (Fig. 4d). Indeed, PML-IV colocalized with TRIM24-TebDB at virtually all telomeres. By co-immunoprecipitation, an interaction between TRIM24 and HA-PML-IV was detected (Extended Data Fig. 8b). Not only was PML recruited to ALT telomeres by TRIM24-TebDB, but several other APB constituents including NBS1, MRE11, and BLM were also highly localized to ALT telomeres (Fig. 4d), implicating TRIM24 in the mechanism that establishes bona fide APBs.

To further understand the functional nature of TRIM24 and PML interactions in ALT, we first depleted PML, which did not affect telomere-localization of endogenous TRIM24 (Fig. 4e and Extended Data Fig. 8c). Consistent with previous observations^43^, tethering PML IV to telomeres significantly increased the size of telomeric foci, marked by GFP-tagged TRF1, in U2OS cells (Fig. 4f). However, tethering of PML to telomeres failed to induce enlarged telomere foci in TRIM24-depleted U2OS cells (Fig. 4f), revealing that TRIM24 may be required for telomere clustering within APBs. To decipher how TRIM24 recruits PML-IV to telomeres, we expressed several TRIM24-TebDB deletion mutants (Fig. 4g). Only the deletion of the coiled-coil domain (CC) of TRIM24 abolished PML-IV telomere recruitment, despite comparable levels of transgene expression (Fig. 4g-h; Extended Data Fig. 8d). These results show that the CC domain of TRIM24 plays a critical role in APB formation.

### SUMOylation of TRIM24 promotes the formation of functional APBs

SUMOylation is a major driver of APB formation^20,44,50^. Western blot analysis of GFP-TRIM24 readily detect several slowly migrating TRIM24 species (Fig. 5a), which have previously been characterized as SUMOylated TRIM24^45^. Bands representing SUMOylated TRIM24 were absent in TRIM24 lacking its PHD or BRD (TRIM24ΔPHD or TRIM24ΔBRD, respectively), as well as in SUMO-Defective TRIM24 (TRIM24-SD); with lysine to arginine mutations at the major SUMOylation sites K723R and K741R^45^ (Fig. 5a). Interestingly, the levels of SUMO-modified chromatin-bound TRIM24 were enhanced following telomere-specific damage inflicted by TRF1-FokI in U2OS cells, which was enhanced after treating TRF1-FokI expressing cells with the broad-HDAC inhibitor Trichostatin A (TSA) (Extended Data Fig. 8e). Thus, these results support the involvement of both SUMOylation and acetylation in regulating TRIM24 telomere localization and subsequent function in ALT.

**Fig. 5:**
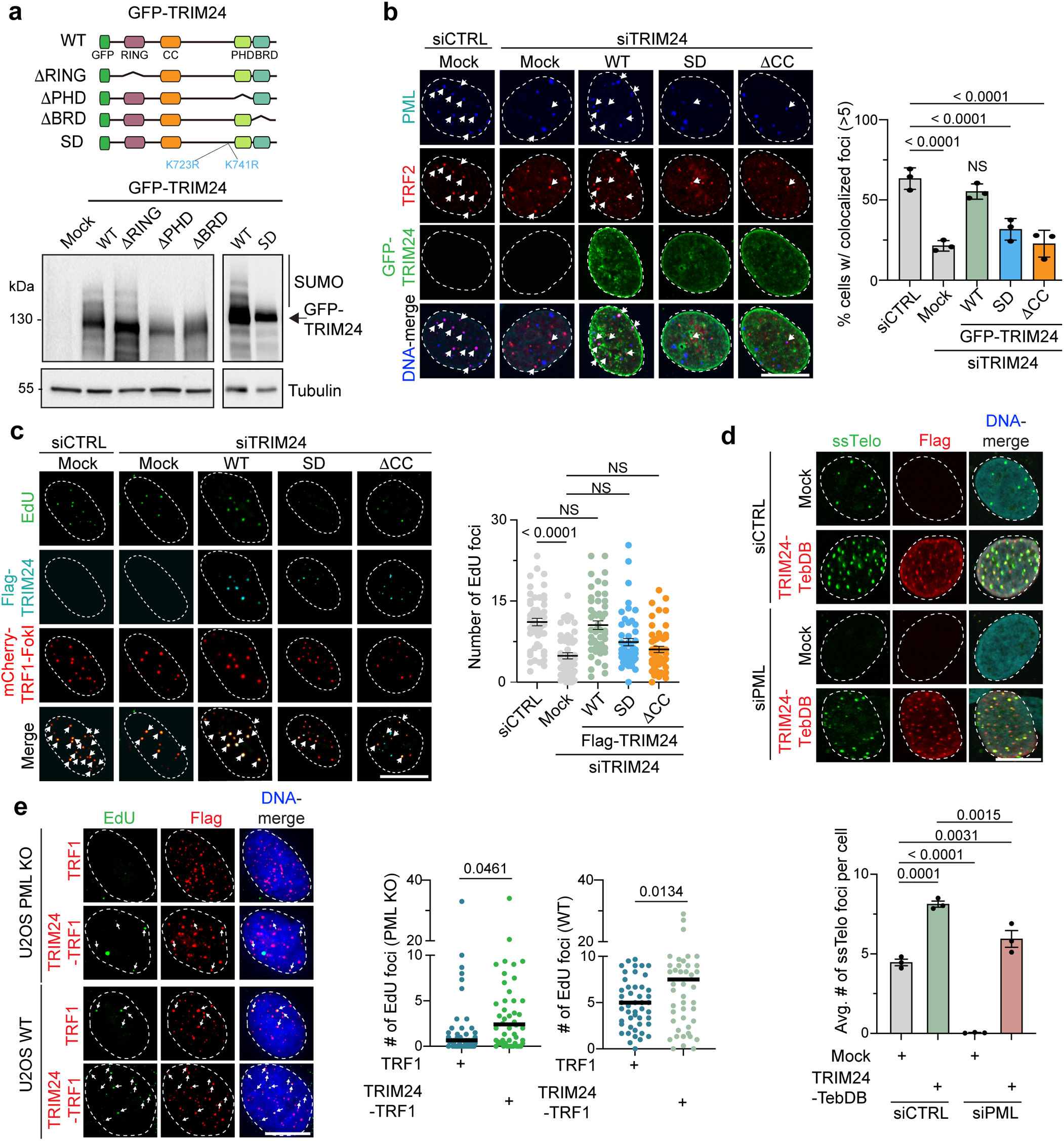
Analysis of TRIM24 in PML dependent and independent ALT. **a,** Depiction of GFP tagged WT TRIM24 and mutants is shown on the top. SUMOylation deficient (SD) TRIM24 has K to R mutations in amino acids 723 and 741. Western blot of lysates expressing WT TRIM24 and derivatives using anti-GFP and Tubulin as loading control is shown. Slower migrating SUMOylated forms of TRIM24 are indicated. **b,** APBs were determined in U2OS cells either treated siCTRL or siTRIM24 3’ UTR followed by complementation with the indicated TRIM24 constructs. Colocalization of PML and telomeres was analyzed by by IF and confocal microscopy. TRF2 indicates telomeres. Cells were treated with 9 μM of RO3306 for 16 hrs prior to analysis to enrich cells in G2. Quantification is shown on the right, n=3, >100. **c,** Nascent telomere DNA synthesis was determined in U2OS cells either treated siCTRL or siTRIM24 3’ UTR followed by complementation with the indicated TRIM24 constructs. TRF1-FokI marks telomeres. Cells were treated with 9 μM of RO3306 for 16 hrs. Quantification is on the right, n=4, >180. **d,** ssTelo foci were analyzed by FISH, IF and confocal microscopy in mock-treated or Flag-TRIM24-TebDB expressing U2OS cells following PML depletion using siRNAs. Quantification is provided below, n=3, >100. **e,** Nascent telomere DNA synthesis in parental or PML KO U2OS cells expressing Flag-TRF1 or Flag-TRIM24-TRF1 was determined. Quantification with two-tailed unpaired T-Test is on the right, n=3, >125. Unless otherwise indicated, statistical analyses were performed with one-way ANOVA, error bars in graphs represent mean ± s.e.m., with P values indicated in the graphs. NS indicates no statistical significance. Dots on the graphs indicates independent experiments. N represents number of biologically independent experiments and indicated number represents total number of cells, or treats analyzed per each experimental group. All scale bars in IF images, 10 μm.

To dissect the mechanistic connection between SUMO and TRIM24 during ALT, we treated U2OS cells expressing TRIM24-TebDB with TAK-981, a potent inhibitor of SUMO-activating enzyme subunit 2 (SAE2) ^51^ and found that it abrogated the recruitment of PML to TRIM24-TebDB marked telomeres (Extended Data Fig. 8f). To directly investigate the role of TRIM24 SUMOylation in APB formation, we examined the abundance of telomere-localized PML in cells lacking endogenous TRIM24 and complemented with various TRIM24 derivatives (Extended Data Fig. 8g). As before, APB formation was reduced upon TRIM24-depletion (i.e. TRF2 and PML colocalization; Fig. 5b) in G2-arrested U2OS cells. Complementation of TRIM24-depleted cells with WT TRIM24 restored the reduced abundance of APBs observed in TRIM24-depleted cells (Fig. 5b). However, expressing TRIM24ΔCC or TRIM24-SD failed to recover telomere-localized PML in TRIM24-deficient cells (Fig. 5b). Importantly, expression of TRIM24 WT but not TRIM24ΔCC or SD mutants rescued the DNA synthesis defects observed in TRIM24-depleted cells (Fig. 5c). Taken together, these findings demonstrate that TRIM24 SUMOylation, as well as its coiled-coil domain, are required to facilitate ALT through an ability to form functional APBs.

### Telomere-tethered TRIM24 promotes ALT independent of PML

We wanted to test further the functional consequences of TRIM24-PML interactions in supporting ALT. Consistent with previous reports^20^, depletion or knockout (KO) of PML in U2OS cells attenuated ALT, as evidenced by the near absence of ssTelo signals and nascent telomere synthesis (Fig. 5d-e and Extended Fig. 9a). Strikingly, using two independent mechanisms of telomere tethering, TebDB or the widely used TRF1 protein, TRIM24 when tethered to telomeres can restore ALT activities including telomere synthesis and telomeric ssDNA formation in PML deficient U2OS cells (Fig. 5d-e and Extended Fig. 9a). Tethering the BTR complex to telomeres has been shown to induce ALT in PML KO cells^20^. The essential function of BLM for ALT is partly by establishing functional APBs^20,43,44^. Since TRIM24-TebDB induced ssTelo and telomere DNA synthesis in PML KO cells, we hypothesized that TRIM24-TebDB might promote PML-independent APB formation by directing the localization of BLM to telomeres in PML-deficient cells. Indeed, TRIM24-TebDB effectively recruited BLM to telomeres in both control and PML-depleted cells (Extended Data Fig. 9b). Conversely, TRIM24 depletion reduced BLM localization at telomeres in G2-arrested U2OS cells (Extended Data Fig. 9c). This was remarkable since PML is considered indispensable for recruiting BLM to telomeres in ALT cancer cells^20^, supporting the role of TRIM24 in functional APB formation.

We next examined the role of BLM in PML-independent ALT induced by TRIM24-TebDB. Depletion of BLM abolished the ssTelo formation induced by TRIM24-TebDB in PML-depleted cells (Extended Data Fig. 9d-e). This result shows that tethering TRIM24 to telomeres does not negate the essential requirement for BLM in ALT, unlike PML. Interestingly, treatment with the SUMOylation inhibitor TAK-981 abolished BLM recruitment to TRIM24-TebDB (Extended Data Fig. 9b), indicating that BLM recruitment by TRIM24-TebDB is SUMOylation-dependent. To further investigate this, we examined the ssTelo formation in PML-depleted cells expressing TebDB fused TRIM24-SD-TebDB. TRIM24-SD-TebDB expressed in PML-depleted U2OS cells reduced ssTelo foci formation compared to TRIM24-TebDB (Fig. 6a), highlighting the functional dependency of TRIM24 SUMOylation in promoting ALT in cells lacking PML. Collectively, these results demonstrate that artificially tethering TRIM24 to telomeres can induce ALT in a PML-independent but BLM- and SUMOylation-dependent manner, likely through the ability of TRIM24 to recruit BLM to ALT telomeres.

**Fig. 6:**
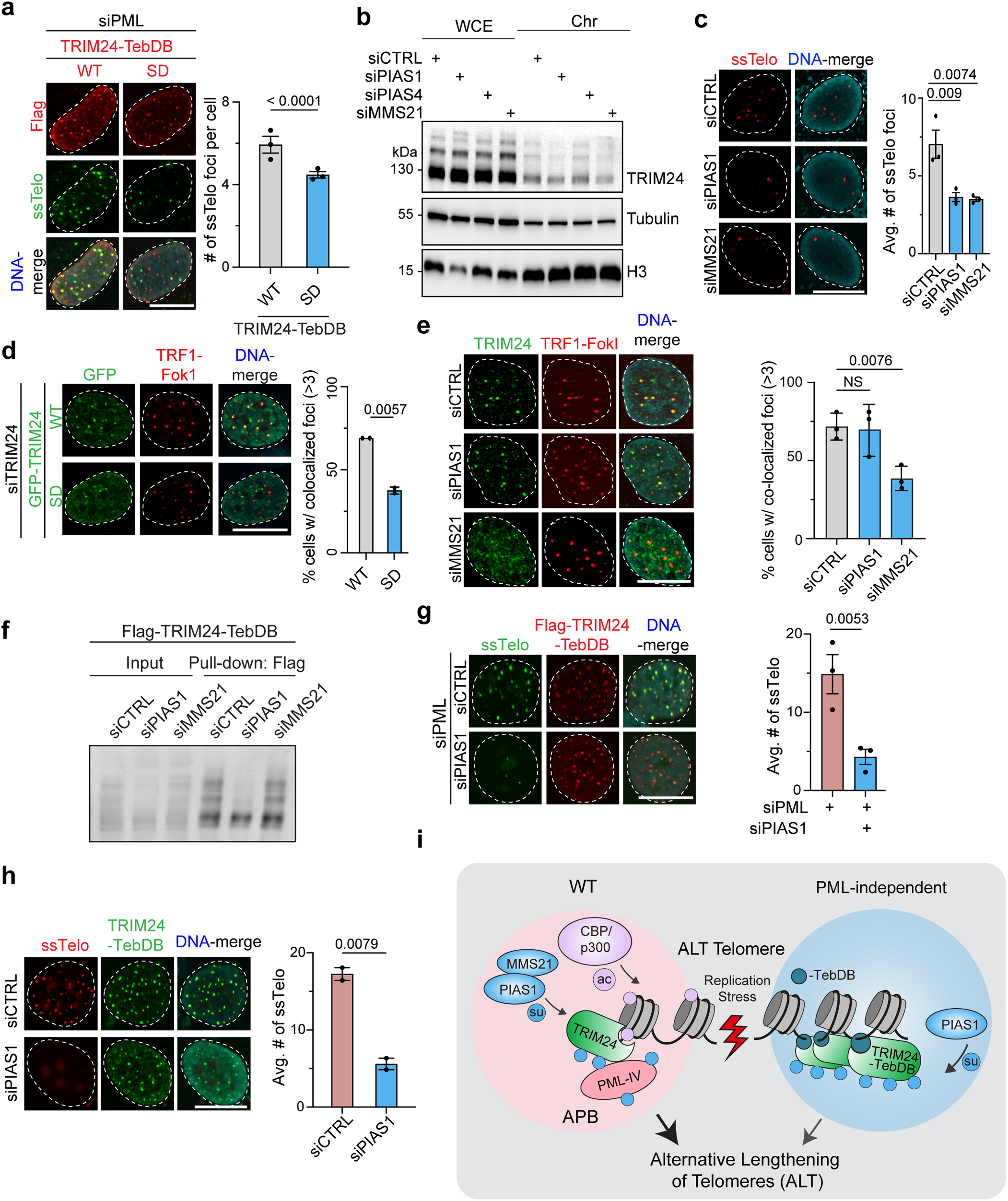
The SUMO E3 ligases MMS21 and PIAS1 regulate TRIM24 to promote ALT. **a,** ssTelo foci were determine by FISH, IF and confocal microscopy in WT or SD TRIM24-TebDB expressing U2OS cells following PML depletion using siRNAs. Quantification is provided on the right, n=3, > 95. **b,** Western blotting of WCE (whole cell extract) and Chr (chromatin fraction) from U2OS cells treated with the indicated siRNAs was performed to detect SUMOylation of endogenous TRIM24. Tubulin and H3 act as loading controls and indicate chromatin fraction with reduced Tubulin and increased H3 signals compared to WCE. **c,** ssTelo foci were analyzed as in **a** following siRNA treatments targeting CTRL, PIAS1 or MMS21. Quantification with one-way ANOVA is on the right, n=3, >200. **d,** Telomere localization of TRIM24-depleted U2OS cells complemented with GFP-tagged TRIM24 WT and SD were analyzed. TRF1-FokI indicates telomeres. Quantification is on the right, n=2, >60. **e,** Localization of endogenous TRIM24 to telomeres in U2OS cells following PIAS1 or MMS21 depletion by siRNAs was determined by IF and confocal microscopy. TRF1-FokI marks telomeres. Quantification with one-way ANOVA is on the right, n=3, > 110. **f,** Western blotting of lysates from WCE and Flag purified immunocomplexes with anti-Flag to detect SUMOylation of Flag-TRIM24-TebDB in U2OS cells following PIAS1 or MMS21 depletion by siRNAs. **g,** ssTelo foci analysis by FISH, IF and confocal microscopy in TRIM24-TebDB expressing PML-depleted U2OS cells following PIAS1 depletion. Quantification is on the right, n=3 >75. **h**, ssTelo foci analysis by FISH, IF and confocal microscopy in TRIM24-TebDB expressing parental U2OS cells following PIAS1 depletion. Quantification is on the right, n=2, > 65. **i,** Summary model for TRIM24 regulation and function in ALT. TRIM24 promotes both PML-dependent APBs and ALT-associated PML-independent bodies to facilitate telomere maintenance through ALT. In APBs, CBP and p300 acetylate H3K23ac to direct TRIM24 to telomeres in a SUMOylation dependent pathway involving MMS21. Once localized to ALT telomeres, TRIM24 undergoes PIAS1-dependent SUMOylation leading to ALT in both APB and APIB forming cells. Unless otherwise indicated, statistical analyses were performed with unpaired two-tailed T-test, error bars in graphs represent mean ± s.e.m., with P values indicated in the graphs. NS indicates no statistical significance. Dots on the graphs indicates independent experiments. N represents number of biologically independent experiments and indicated number represents total number of cells, or treats analyzed per each experimental group. All scale bars in IF images, 10 μm.

### TRIM24 collaborates with MMS21 and PIAS1 to promote ALT

Given our identification of TRIM24 SUMOylation in ALT, we sought to identify the SUMO E3 ligase(s) involved in this pathway, focusing on PIAS1, PIAS4 and MMS21 due to their established roles in the DDR^52^ and ALT^53^. First, we observed that the SUMOylation of chromatin-bound TRIM24 was reduced upon the depletion of either PIAS1 or MMS21, but not PIAS4 (Fig. 6b; siRNA-depletion of proteins shown in Extended Data Fig. 10a). Depletion of PIAS1 or MMS21 led to a noticeable decrease in telomeric ssDNAs (Fig. 6c). We speculated that SUMOylation of TRIM24 might regulate its recruitment and/or retention at telomeres. In contrast to WT TRIM24, SUMO-defective TRIM24-SD localization to telomeres was markedly reduced (Fig. 6d). Furthermore, depletion of MMS21 significantly reduced the telomere localization of TRIM24, while depletion of PIAS1 had little to no effect (Fig. 6d). Of note, upon TRIM24 tethering to telomeres, PIAS1-depletion led to reduced TRIM24 SUMOylation while MMS21-deficiency had no effect (Fig. 6f). These data indicate that once TRIM24 is localized to telomeres, MMS21 is dispensable for TRIM24 SUMOylation, which becomes dependent on PIAS1. In line with the involvement of PIAS1 in mediating TRIM24 SUMOylation at telomeres, expression of TRIM24-TebDB robustly recruited PIAS1 but not MMS21 to these sites, which was dependent on the CC domain of TRIM24 (Extended Data Fig. 10c-d). Functionally, we observed a substantial reduction in ssTelo foci in TRIM24-TebDB expressing cells that were either PML proficient or deficient (Fig. 6g-h), indicating the importance of PIAS1-mediated SUMOylation in promoting ALT activities. These results define a distinct pathway involving both MMS21- and PIAS1-mediated SUMOylation of TRIM24 in the ALT pathway. MMS21 SUMOylation localizes/recruits TRIM24 to telomeres whereupon PIAS1 interacts with TRIM24 at telomeres through its CC-domain to promote TRIM24 SUMOylation and ALT.

This comprehensive analysis defines a sequence of molecular events involving TRIM24 that preserves telomeres specifically in ALT cancer cells. First, chromatin acetylation by CBP/p300 and SUMOylation by MMS21 localize TRIM24 to telomeres where it connects telomeres to APBs and telomere DNA synthesis (Fig. 6i). These steps rely on PIAS1-mediated SUMOylation and BLM. In the absence of PML, we identified that telomere-tethered TRIM24 can induce PML-independent APB-like bodies that rely on SUMOylation and BLM to facilitate telomere maintenance in ALT cancer cells (Fig. 6i). Crucially, disrupting this sequence events abrogates ALT activity implicating this TRIM24-CBP/p300-SUMO axis in directing critical processes required for telomere length maintenance in ALT cancer cells.

## Discussion

In this study, we developed a novel strategy, BLOCK-ID, to identify proteins specifically associated with obstructed replication forks. BLOCK-ID combines unbiased mass spectrometry and single-cell fluorescence microscopy imaging of protein dynamics at stressed replication forks, allowing orthogonal approaches to study and validate replication stress response factors in cells. BLOCK-ID harnesses an engineered genomic locus containing the Lac Operator (LacO) repeat array that when bound by the Lac Repressor (LacI) forms a protein complex barrier that impedes replication fork progression. While several powerful approaches have been developed to detect replication stress associated proteins, including iPOND and Nascent Chromatin Capture, BLOCK-ID does not require nucleotide labeling, instead enabling direct detection of proteins at stressed replication forks by proximity-dependent biotinylation (Fig. 1a). This powerful feature of BLOCK-ID overcomes limitations in capturing highly dynamic proteins, preserving the capacity to capture proteins that are transiently and/or weakly associated with stressed replication forks. Comparisons of proteins enriched between BLOCK-ID, iPOND and NCC showed similar levels of overlap (Extended Data Fig. 1d-f), showing good concordance between BLOCK-ID and these established methodologies. The utility of BLOCK-ID was validated by our identification of nearly half of the 42 human Bromodomain proteins associated with stressed replication forks (Fig. 1e), highlighting the importance of this chromatin reader protein family in DNA replication responses^31^.

While BLOCK-ID has several advantages as noted above, this technique might not capture the landscape of proteins associated with stressed replication forks throughout the genome in different environments. Additionally, the slow kinetics of BirA*^23^ might favor the identification of proteins associated with collapsed replication forks. However, employing biotin ligases with faster kinetics such as Turbo-ID^54^ in future studies might allow for more temporal investigation of stressed replication forks by BLOCK-ID. Regardless, our study establishes BLOCK-ID as a powerful, broadly applicable new approach to interrogate the replication stress response, whose use will likely deliver new mechanistic insights of how replication forks are protected and processed from stresses that when impaired drive genome instability and cancer initiation.

Here, we focused on TRIM24, which we characterize as a new replication stress response factor with an unexpected but essential role in telomere extension by the ALT mechanism. We determined that TRIM24 localizes to telomeres in ALT cancer cells where it contributes to replicative stress mitigation and telomere structural integrity (Fig. 1g, and Fig. 2f). Furthermore, disruption of TRIM24 robustly interferes with telomere DNA synthesis (Fig. 2d), leading to telomere destabilization and rapid shortening (Fig. 2e). Mechanistically, we identified that p300 and CBP-dependent acetylation of histone H3 at lysine23 (H3K23ac) mediates TRIM24 recruitment to telomeres (Fig. 3a, and Fig. 3e). In agreement, prior structural studies determined that the PHD-BRD reader cassette within TRIM24 preferentially binds to unmodified histone H3 lysine 4 and H3K23ac^33,55^. The PHD-BRD of TRIM24 forms contacts with each other in the crystal structure and mutations in either the PHD or BRD disrupt binding to H3K23ac^33^. Accordingly, TRIM24 mutants lacking these reader modules did not associate with telomeres (Fig. 3a). Remarkably, we found that once recruited this CBP-p300-H3K23ac regulatory network is dispensed with (Fig. 4b and Extended Fig. 8a), becoming no longer necessary to retain TRIM24 at telomeric chromatin. These results suggested that the main function of p300/CBP is to modify chromatin within ALT telomere that signals the recruitment of binding to these genomic loci. However, we cannot rule out that p300/CBP may have additional targets at telomeres that act upstream of TRIM24 recruitment in ALT. For example, TRF2 is acetylated by p300^56^, which may also contribute to TRIM24 accumulation at telomeres and ALT. Future studies will be needed to explore further the acetylome at telomeres that mediate their maintenance in ALT.

In addition to acetylated H3, TRIM24 also binds unmodified H3K4 and methylated H3K9, a main repressive histone mark associated with ‘closed’ heterochromatin, including at stalled replication forks where it protects against nucleases that attack and degrade the fork^7^. TRIM24s ability to bind both methylated and acetylated histones is intriguing as it allows TRIM24 the capacity to bind to chromatin irrespective of chromatin state, including potentially at telomeres. The dual PTM reader function could enable TRIM24 to stably associate at telomeres whereby it is initially recruited via H3K23ac and then retained through H3K9me3 binding. Notably, H3K9me3 at telomeres, catalyzed by SETDB1^57^, which like TRIM24 as demonstrated here, are implicated in establishing and maintaining productive ALT. Interestingly, a potentially related observation was demonstrated for TRIM24 as a transcription cofactor of p53, where TRIM24 specifically acts at tightly closed chromatin regions marked by unmethylated H3K4 in mouse embryonic stem cells^55^. In these cells, TRIM24 simultaneously binds to both p53 and unmethylated H3K4 to promote the opening of condensed chromatin for transcription activation^55^. Thus, multivalent interactions between chromatin and histone marks may govern the function of TRIM24 in diverse biological processes including transcription, and as demonstrated here, ALT maintenance. Future work could dissect TRIM24s chromatin interactions at telomeres specifically, including possible links to SETDB1-dependent H3K9me3, to elucidate the functional contributions to ALT.

In addition to the CBP/p300 network that regulates TRIM24 recruitment to telomeres, the relationship between TRIM24 and ALT-associated PML bodies was particularly striking. We initially found that disrupting TRIM24 impaired APB formation. By artificially tethering TRIM24, through fusion with TebDB or TRF1, we observed the striking *de novo* enrichment of PML protein to all telomeres (Fig. 4c, and 4d). Through deletion studies, we uncovered this activity requires the SUMOylation of TRIM24 by PIAS1 (Fig. 6b, and 6f), which facilitates interactions with other key mediators of ALT such as the BLM helicase (Extended Data Fig. 9b). Prior studies showed that tethering BLM, and its cognate interacting proteins TOP2A and RMI1/2, at telomeres can bypass the PML dependency for ALT induction^20^. While telomere-tethered TRIM24 can recruit BLM to telomeres, we found that TRIM24 depletion resulted in the loss of BLM localization to ALT telomeres (Extended Data Fig. 9c). This positions TRIM24 upstream of BLM in the mechanism of APB formation. Consistent with this notion, TRIM24 depletion leads to inefficient telomere clustering within APBs (Fig. 4f). It is tempting to speculate that SUMOylated TRIM24 may promote the telomere-localization of BLM and PML, both of which bind to SUMO through SIM motifs^44,58^. Therefore, TRIM24 recruitment at telomeres might seed and bridge the formation of APBs onto telomeric chromatin containing SUMOylated TRIM24, which represent necessary conduits for ALT-mediated telomere extension. It is worth noting that TRIM24 was previously implicated in aberrant PML dynamics. The fusion oncoprotein PML-RARα, which forms dysregulated interactions with TRIM24^41^, impairs the proper formation of PML nuclear bodies, leading to the mislocalization of BLM and subsequently defects in homologous recombination^59^. This implicates, along with our data provided here, the role of TRIM24 as a regulatory factor for functional formation of PML nuclear bodies, including at telomere.

Remarkably, we also found that telomere tethering of TRIM24 was sufficient to induce features of ALT activity, most notably *de novo* synthesis of telomeric DNA, in cells devoid of PML (i.e. PML KO cells) (Fig. 5d and 5e). These findings provide several important mechanistic insights into the mechanisms of ALT. For example, TRIM24 deficiency could affect ALT indirectly by altering transcriptional regulation of cells. The ability of TRIM24 to directly promote ALT while tethered to telomeres, including in PML-deficient cells and telomerase-positive cells, strongly argues against this possibility. These results suggest that TRIM24 binding to telomeric chromatin may be sufficient to stimulate ALT, again through BLM and SUMOylation of TRIM24 but independent of PML. How PML-independent ALT telomere DNA synthesis is initiated is not known. Recently, BLM was shown to initiate ALT-telomere DNA synthesis at misprocessed Okazaki fragments^60^. Tethering of SUMO3 using a chemical dimerizer can also promote ALT features by recruiting BLM, RAD51AP1, and RAD52^61^. Among these factors, BLM recruitment to telomeres by SUMO3 induces ALT in PML KO cells in a SUMOylation-dependent manner^61^, This is consistent with our findings that mutation of two key SUMOylation sites on TRIM24 reduced its ability to promote ALT, highlighting again the involvement of SUMOylation in ALT. Determining how TRIM24 can bypass the requirement for PML should be explored further. Identifying SUMO-dependent interactors of TRIM24 specifically in PML-deficient ALT cells at telomers may provide key insights into the molecular mechanisms regulating APIBs formation that we report here. Regardless, the fact that nascent telomere DNA synthesis can be stimulated in the absence of PML by TRIM24 is an important observation, raising further questions about the functional role of both PML and TRIM24.

In summary, we have identified a p300/CBP/TRIM24 chromatin signaling pathway that promotes ALT telomere maintenance via BLM and PIAS1-mediated SUMOylation. Our mechanistic findings highlight a previously uncharacterized role for acetylation signaling in ALT that TRIM24 governs. The finding that TRIM24 recruits PML to telomeric chromatin provides a mechanistic basis for the known telomere localization and DNA synthesis of telomeres within APBs in ALT. The strong reduction in ALT activities, including the rapid loss of telomeres, in TRIM24-deficient cells demonstrates the essential function of this protein in ALT telomere maintenance. Our discoveries have important clinical implications, identifying this new ALT chromatin signaling pathway involving several acetylation signaling proteins, as potential targets in ALT cancers.

## Methods

### Plasmids

To generate mycBirA*-LacI, the LacI sequence was amplified by PCR from the mCherry-LacI vector^30^ and inserted into the mycBirA* pcDNA3.1 vector^23^ using the XhoI and AflII restriction enzyme sites. This construct was then subcloned into the pCW57.1 vector using the NheI and EcoRI restriction enzyme sites. Functional domains of TRIM24 were identified using the UniProt/Prosite database. Deletion mutants of TRIM24 were generated by PCR amplification and inserted into custom-built expression vectors with N-terminal eGFP (GFP-pcDNA3.1) or Flag (Flag-pcDNA5 FRT/TO) tags. SUMO-deficient TRIM24 mutants were generated via site-directed mutagenesis and inserted into both GFP-pcDNA3.1 and Flag-pcDNA5 FRT/TO vectors using the XhoI and AlfII restriction enzyme sites. The DNA binding domain of Teb1 was amplified by PCR and C-terminally fused to TRIM24 and PML IV through overlap extension PCR. The resulting PCR products were inserted into GFP-pcDNA3.1 and Flag-pcDNA5 FRT/TO vectors using the XhoI and AlfII restriction enzyme sites. TRIM24 shRNAs (C1 and C2) were cloned into the pLKO.1 TRC vector according to the protocols from Addgene.

### Cell lines and cell line generation

All cells used in this study were cultured in Dulbecco‘s modified Eagle‘s medium (DMEM) supplemented with 10% fetal bovine serum (FBS), 2 mM L-glutamine, 100 U/ml penicillin, and 100 µg/ml streptomycin, and maintained at 37°C in a 5% CO₂ atmosphere. For non-dividing conditions, cells were cultured in DMEM supplemented with 0.1% FBS, 2 mM L-glutamine, 100 U/ml penicillin, and 100 µg/ml streptomycin, and maintained at 37°C in 5% CO₂ for two days. mycBirA*-LacI construct cloned into the pCW57.1 vector and shRNAs cloned into the pLKO.1 vector were introduced into target cells via lentiviral transduction. mycBirA*-LacI or shRNAs, along with the packaging vectors pPAX2 and pMD2.G, were co-transfected into HEK293T cells using Lipofectamine 2000 (ThermoFisher) according to the manufacturer’s instructions. Lentivirus particles were harvested at 48 and 72 h post-transfection, centrifuged at 500 xg for 10 min, and filtered through 0.45 µm PES syringe filters to remove cell debris. The virus-containing media were then applied to the target cells. The following day, the lentivirus media was replaced with fresh culture media containing the indicated antibiotics for selection. Detailed information on the shRNAs used in this study is provided in Supplementary Table 2.

### Immunofluorescence

Cells were plated on glass coverslips and subjected to the desired experimental conditions. Prior to fixation, a pre-extraction step was performed by incubating the cells on ice do 5 min in CSK buffer containing 10 mM PIPES, pH 6.8, 100 mM NaCl, 300 mM sucrose, 3 mM MgCl₂, 1 mM EGTA and incubated on ice for 5 min. The cells were then washed three times with cold PBS and fixed with 3% paraformaldehyde (PFA). Following fixation, cells were permeabilized with 0.4% Triton X-100 for 15 minutes at room temperature. Blocking was performed for 1 h at room temperature using a buffer containing 3% BSA and 0.4% Triton X-100. Then, primary antibodies, diluted in the blocking buffer, were applied and incubated overnight at 4°C. The next day, Alexa Fluor-conjugated secondary antibodies (ThermoFisher) were applied and incubated for 1 h at room temperature. Then, coverslips were mounted onto slides with Vectashield mounting medium containing DAPI (Vector Laboratories). Immunofluorescence signals were visualized using an inverted FV3000 scanning confocal microscope (Olympus) with Olympus PlanApo N 60x/1.40 Oil 0.17/FN22 Microscope Objective (Olympus), controlled by FW31S software (Olympus). Detailed information on the antibodies used in this study is provided in Supplementary Table 3.

### qPCR

Total RNA was extracted from cells using the RNeasy kit (Qiagen) as instructed by the manufacturer. cDNA was generated from the extracted RNA using Superscript III First-Strand Synthesis kit (ThermoFisher). Quantitative PCR was conducted on the StepOnePlus Real-time PCR system (Applied Biosystems) with Fast SYBR™ green master mix (ThermoFisher). Detailed information on the qPCR primers in this study is provided in Supplementary Table 4.

### Clonogenic assays

siRNAs were introduced using RNAiMAX (ThermoFisher) following the manufacturer’s recommendations. Next day, cells were trypsinized and seeded at a density of 500 cells per well in triplicate in six-well plates. Cells were incubated for 10 to 14 days until colonies formed. Colonies were stained with 0.5% crystal violet in 20% ethanol and manually counted. The efficiency of siRNA-mediated knockdown was assessed by Western blot or qPCR. Detailed information on the siRNAs used in this study is provided in Supplementary Table 2.

### Cell cycle analysis

Cell cycle analysis was performed as previously described^62^, with minor modifications. Briefly, cells were incubated in medium containing 10 µM BrdU for 30 min prior to collection. After trypsinization, cells were washed twice with PBS and fixed in 70% ethanol for 30 min on ice. Fixed cells were then permeabilized using a buffer containing 2M HCl and 0.5% Triton X-100 in PBS for 30 min at room temperature on a rocking platform. Next, cells were incubated with 0.1 M Na₂B₄O₇ (pH 8.5) and washed with 1% BSA/PBS containing 0.3 mM EDTA. Cells were then incubated with an anti-BrdU antibody for 1 h, followed by Alexa Fluor-conjugated secondary antibodies for 45 min at room temperature. After washing again with 1% BSA/PBS containing 0.3 mM EDTA, cells were stained with 4 µg/ml propidium iodide and treated with 100 µg/ml RNase A for 30 min. Samples were analyzed by BD LSRFortessa SORP Flow Cytometer (BD Biosciences), and data were processed using FlowJo software.

### Detection of ssTelo foci

Detection of ssTelo foci was carried out as previously described^20^. After completing the immunofluorescence procedure as described earlier, cells were re-fixed with 3% PFA. Then, the coverslips were incubated with the blocking buffer containing 500 µg/mL Rnase A for 1 h at 37°C. After Rnase A treatment, the coverslips were dehydrated through a series of ethanol washes (70%, 90%, and 100%). Once air-dried, FITC-OO[TTAGGG]3-labeled PNA probe (PNA Bio), diluted in 10 mM Tris pH 7.4, 70% formaldehyde, 1 mg/ml FISH blocking reagent, were applied and incubated with the coverslips at room temperature for 3 h. The coverslips were then washed twice with 70% formaldehyde in PBS, followed by two additional washed with 1X PBS. The coverslips were mounted onto slides with Vectashield mounting medium containing DAPI.

### Immunoprecipitation and Western blotting

Cells were lysed using 1 ml of NETN buffer supplemented with 1X HALT protease inhibitor and 1X HALT phosphatase inhibitor. To detect SUMOylation of TRIM24 and TRIM24-TebDB, cells were lysed using 1 ml of RIPA buffer (50 mM TRIS pH 7.0, 150 mM NaCl, 1 % NP-40, 1 % Sodium deoxycholate, 0.1 % SDS). After incubation with 1 unit of Turbonuclease (Accelagen), insoluble components were removed by centrifugation at 16,500 × g for 10 min at 4°C. The supernatant was collected and incubated with 1 µg of the desired antibodies and 20 µl of protein A beads (ThermoFisher) overnight at 4°C. The following day, beads were washed three times with the lysis buffer and then with 50 mM TRIS three times. Bound proteins were eluted from the beads by adding 2X SDS-PAGE sample buffer. The eluates were boiled at 95°C for 5 min, separated by SDS-PAGE, and transferred onto a nitrocellulose membrane for 90 min at 110 V. After blocking with 5% BSA in TBS-T (0.05% Tween-20), primary antibodies, diluted in the blocking buffer, were applied overnight at 4°C. The membrane was washed three times with TBS-T for 10 min each at room temperature and then incubated with HRP-conjugated secondary antibodies (Cell Signaling Technologies) for 1 h at room temperature. Chemiluminescence signals, following application of ECL (GE Healthcare), were detected using Chemidoc MP Imaging System (Bio-Rad).

### SIRF assay

Cells on coverslips were incubated with 125 mM of EdU for 8 min and then treated with 4 mM HU for 3 h. After fixation with 3 % PFA, EdU was conjugated to biotin by incubating click reaction buffer containing 1X PBS, 2mM copper sulfate, 10 μM biotin-azide and 100 mM Sodium ascorbate. Subsequently, primary antibodies against biotin and either TRIM24 or BLM were applied and incubated at 4°C overnight. The next day, PLA was performed using Duolink® PLA kit (Sigma Aldrich) according to the manufacturer’s instruction. Briefly, after overnight incubation with the primary antibodies, In Situ PLA probes (anti-mouse and anti-rabbit) were applied to coverslips for 1 h at 37 4°C. The PLA ligase mix was then added and incubated for 30 minutes at 37°C. Following three washes, the PLA amplification mix was applied and incubated for 90 minutes at 37°C. After additional washes, the coverslips were mounted on slides using Vectashield mounting medium containing DAPI.

### Telomere DNA synthesis detection using EdU

For the siRNA-mediated knockdown conditions, 5 pM of siRNAs was introduced to 800,000 cells using Dharmafect (Horizon Discovery). 48 h later, cells were transfected with a FLAG-tagged TRF1-FokI expressing plasmid^12^. Cells were pulsed with 10μM of EdU for 1 h before fixation. Cells on coverslips were washed with PBS and fixed by incubating with 2% PFA for 10 min. Cells were permeabilized with 0.1% (w/v) sodium citrate and 0.1 % (v/v) Triton X-100 for 5 min and then washed with PBS before incubating with blocking solution (1mg/mL BSA, 10% normal goat serum, 0.1% Tween) for 30 min. Primary antibodies were diluted in blocking solution and added to cells overnight at 4°C. Next, cells were washed three times with PBS for 5 min and incubated with Alexa Fluor-conjugated secondary antibodies (Life Technologies) for 1h at room temperature. After incubation, cells were washed three times, fixed again in 2% PFA for 10 min followed by two additional PBS washes prior to performing click-reaction using the Click-IT Plus EdU Cell Proliferation Kit (Invitrogen) to detect EdU. For experiments with A-485 and SGC-CBP30, U2OS cells inducibly expressing mCherry-TRF1-FokI were cultured with inhibitors at the indicated concentrations for 3 days. 24 h prior to harvest, cells were induced with doxycycline (40ng/μL1) followed by the addition of 1μM of tamoxifen (4-OHT, Sigma) and 1μM of Shield1 Ligand (Takara Clontech) for 3 h before harvest. Cells were pulsed with EdU (10μM) for 1 h before processing as above. For the complementation experiments, 48 h post knockdown using siCTRL or TRIM24 3’UTR siRNA, cells were transfected for 6 h with mCherry-tagged TRF1-FokI expressing plasmid alone or in combination with FLAG-tagged variants of TRIM24. Next day, cells were pulsed with EdU (10μM) for 1hr and then harvested for IF as described above.

### Telomere restriction fragment analysis by pulsed-field gel Electrophoresis (PFGE)

Telomere gels were performed using telomere restriction fragment (TRF) analysis. Genomic DNA was extracted from the different cell lines and digested overnight with AluI and MboI restriction enzymes (NEB). Digested DNA was purified and 2–5 μg of DNA was run on a 1% PFGE agarose gel (Bio-Rad) in 0.5 × TBE buffer using the CHEF-DRII system (Bio-Rad) at 6V cm^−1^; initial switch time 1 s, final switch time 6 s, for 17 h at 14°C. The gel was then dried for 2 h at 60°C, denatured in a 0.5 N NaOH 1.5 M NaCl solution, and neutralized. Gel was hybridized with ^32^P-labeled (TTAGGG)4 oligonucleotides overnight at 55°C in UltaHyb Hybridization Buffer (ThermoFisher Catalog number: AM8669). The next day, the membrane was washed three times in 2x SSC and once in 2x SSC supplemented with 0.5% SDS, exposed onto a phosphor screen, and scanned using Typhoon 9400 PhosphoImager (GE Healthcare).

### Chromosome Orientation FISH (CO-FISH)

CO-FISH was performed as described^63^. In brief, cell cultures were incubated with 7.5mM BrdU and 2.5mM BrdC for ∼14 h. After removal of nucleotide analogs, colcemid (GIBCO) was added for ∼2h and cells were harvested by trypsinization, swelled in 0.075M KCl for approximately 7 min at 37°C and fixed in cold fixative (70% Methanol: 30% Glacial Acetic Acid). Metaphase chromosomes were spread by dropping them onto slides, then RNase A (0.5 mg/mL) was treated. Slides were incubated in 2 × SSC containing 0.5 mg/mL Hoechst 33258 for 15 mins in the dark and irradiated for 3 min (5.4 × 10^5^ J/m^2^, energy 5400) in a UV Stratalinker 2400 (Stratagene). The nicked BrdU/C substituted DNA strands are degraded by Exonuclease III digestion for 1 h. The slides were then washed in PBS, dehydrated by ethanol washes and allowed to air dry completely. The remaining strands were hybridized with fluorescence labeled DNA probes. The positive telomere strand (polymerized by lagging strand synthesis) was labeled using Alexa Fluor 488-conjugated (TTAGGG)4, and the negative telomere strand (polymerized by leading strand synthesis) was labeled using Alexa Fluor 568 conjugated (CCCTAA)4. Prior to hybridization of the first PNA, DNA is denatured by heating at 70°C for 10 min, and then incubated for 2 h at room temperature. Slides were washed for 15 min with Wash Solution A (70% Formamide and 10mM Tris-HCl pH 7.0-7.5, BSA), dried, and then incubated with the second PNA for 2 h at room temperature. The slides were then washed again twice for 15 min with Wash Solution A and 3 times with Wash Solution B (0.1M Tris-HCl pH7.2, 0.15M NaCl and 0.08% Tween) for 5 min at room temperature. Finally, cells were dehydrated by ethanol washing as mentioned above and mounted using ProLong Gold Mounting Media with DAPI. Metaphase chromosomes were visualized by conventional florescence microscope with a 63X Plan λ objective (1.4 oil) on a Nikon 90i microscope.

### BioID pull-down and on-bead digestion

BioID pull-down was performed as previously described^23,64^. Briefly, biotinylation in living cells was achieved by adding 50 μM biotin to the culture media and incubating overnight. The following day, 2 × 10^8^ cells from each experimental set were counted and lysed using BioID lysis buffer (50 mM Tris, pH 7.4, 500 mM NaCl, 0.4% SDS, 1 mM dithiothreitol, and 1X HALT protease inhibitor). After lysis, the cell suspension was subjected to two rounds of tip sonication at power 1 for 1 min each (20 seconds on, 20 seconds off) using a FB705 ThermoFisher sonicator. Insoluble cellular debris was removed by centrifugation at 16,500 xg for 10 min at 4°C. The supernatant was then incubated with Dynabeads MyOne C1 streptavidin beads (ThermoFisher) overnight at 4°C with gentle rotation. After incubation, the beads were collected using a magnetic stand and sequentially washed once with Wash buffer 1 (2% SDS), once with Wash buffer 2 (0.1% deoxycholate, 1% Triton X-100, 500 mM NaCl, 1 mM EDTA, and 50 mM HEPES), once with Wash buffer 3 (250 mM LiCl, 0.5% NP-40, 0.5% deoxycholate, 1 mM EDTA, and 10 mM Tris, pH 7.4), and once with 50 mM Tris. The captured proteins were subjected to on-bead digestion as previously described^65^. Briefly, the beads were resuspended in a solution containing 50mM triethylammonium bicarbonate (TEABC, Sigma-Aldrich) and 2 M urea. Reducing of proteins was performed by incubating the beads with 20 mM DTT for 30 min, followed by alkylation with 20 mM iodoacetamide for 30 min in the dark. Then, 1 μg of sequencing grade trypsin (Promega) was added to the beads, and the mixture was incubated at 37°C overnight on a shaker. The following day, the digested peptides were collected and cleaned using a custom-built reversed-phase C18 column.

### Liquid chromatography-mass spectrometry (LC-MS/MS) analysis

The LC-MS/MS analysis was conducted as described in the previous publications with minor modifications^66,67^. The peptides were sequenced using the Orbitrap Fusion Lumos Tribrid Mass Spectrometer and the Ultimate3000 RSLCnano liquid chromatography system (ThermoFisher). The peptides, which were reconstituted in 15 μl of 0.1% formic acid, were injected into an Acclaim PepMap100 Nano-Trap Column (100 μm × 2 cm, ThermoFisher Scientific, San Jose, CA, USA) containing 5 μm C18 particles at a flow rate of 5 μl per min. A linear gradient of 8% to 28% solvent B (0.1% formic acid in 95% acetonitrile) was used to separate the peptides at a flow rate of 300 nl/min over 95 min on an EASY-Spray column (50 cm x 75 µm ID, ThermoFisher) packed with PepMap RSLC C18 and 2 µm C18 particles (ThermoFisher). An EASY-Spray ion source was used for ionization at 2.4 kV. The mass spectrometry analysis was conducted in a data-dependent mode with a full scan in the mass-to-charge ratio (m/z) range of 350 to 1800 in the “Top Speed” setting, with a three-second cycle. MS1 for the precursor ions and MS2 for the fragmentation ions were acquired at a resolution of 120,000 and 30,000 at an m/z of 200, respectively. The MS2 fragmentation was carried out using the higher-energy collisional dissociation (HCD) method at 32 of normalized collision energy (NCE) with 5% stepped NCE. The automatic gain controls were set to 1 million ions for MS1 and 0.05 million ions for MS2, and the maximum ion injection time was set to 50 milliseconds for MS1 and 100 milliseconds for MS2. The precursor isolation window was set to 1.6 m/z with a 0.4 m/z of offset, and dynamic exclusion was set to 30 seconds while singly charged ions were rejected. Calibration was carried out using the lock mass option (m/z 445.1200025) from ambient air.

### Database search

The database search was conducted as described in the previous publications with minor modifications ^66,67^. To identify and quantify proteins, the acquired spectra were searched against the human UniProt database (released in December 2019, containing protein entries with common contaminants) using the SEQUEST search algorithm embedded in the Thermo Proteome Discoverer platform (version 2.2, ThermoFisher Scientific). For MS/MS preprocessing, the top 10 peaks in each 100 m/z window were selected for the database searches. The following search parameters were employed: a) trypsin was set as a proteolytic enzyme (with up to two missed cleavages); b) 20 ppm was set as peptide mass error tolerance; c) 0.02 Da was set as fragment mass error tolerance; d) carbamidomethylation of cysteine (+57.02146 Da) was set as fixed modification; e) oxidation of methionine (+15.99492 Da) and protein acetylation (+ 42.01056 Da) on N-terminus were set as dynamic modifications; and f) the minimum peptide length was set to 6 amino acids. Peptides and proteins were filtered at a 1 % false-discovery rate (FDR) at the PSM level using the percolator node and at the protein level using the protein FDR validator node, respectively. For BLOCK-ID candidate identification, proteins identified by BirA* were excluded. Additionally, unique proteins found in the proliferating condition and those enriched more than 1.5-fold compared to the growth-arrested condition were included.

### Quantification and statistical analysis

Statistical calculations were done using Prism (v. 10.2.0). Two-tailed Student’s t-test, one-way analysis of variance by a Dunnett multiple comparison test and Mann-Whitney test were used to determine statistical significance as indicated in each figure legend. Image analysis was done using ImageJ software (v. 1.53a).

## Acknowledgement

We thank Dr. Roger Greenberg (University of Pennsylvania) for generously providing the U2OS 256X cells. We grateful to Dr. Eros Denchi (National Cancer Institute) for insightful feedback on the manuscript. We thank all members of the Miller and O’Sullivan laboratories for valuable discussions. For this study, the Miller laboratory is supported by NCI (RO1CA198279, and CA250905), and Cancer Prevention and Research Institute of Texas (RP220330); the O’Sullivan laboratory is supported by NCI (RO1CA262316, RO1CA262316, and P30CA047904); and the Na laboratory is supported by NIH (S10OD021844).

## Contributions

D.K. and K.M.M. conceived the study. D.K. designed and performed the experiments unless otherwise indicated. R.B. performed and analyzed the PFGE, APB, telomere synthesis, telomere metaphase, and telomere CO-FISH assays. R.W.B. performed and analyzed telomere metaphase experiments. S.W. conducted the screening of HATs and the p300/CBP-dependent telomere localization of TRIM24. D.L. performed the cell cycle assay. D.L. and R.P. screened the localization of BRD proteins at BLOCK-ID foci and assessed HU sensitivity. S.T.O. and C.H.N. developed and implemented the analytical mass-spectrometry pipeline for BLOCK-ID. K.M.M. and R.J.O. supervised the study and provided funding. D.K. and K.M.M. wrote the initial manuscript with input from all other authors.

## Supplementary Information

### Extended Data Figure Legends

**Extended Data Fig. 1:**
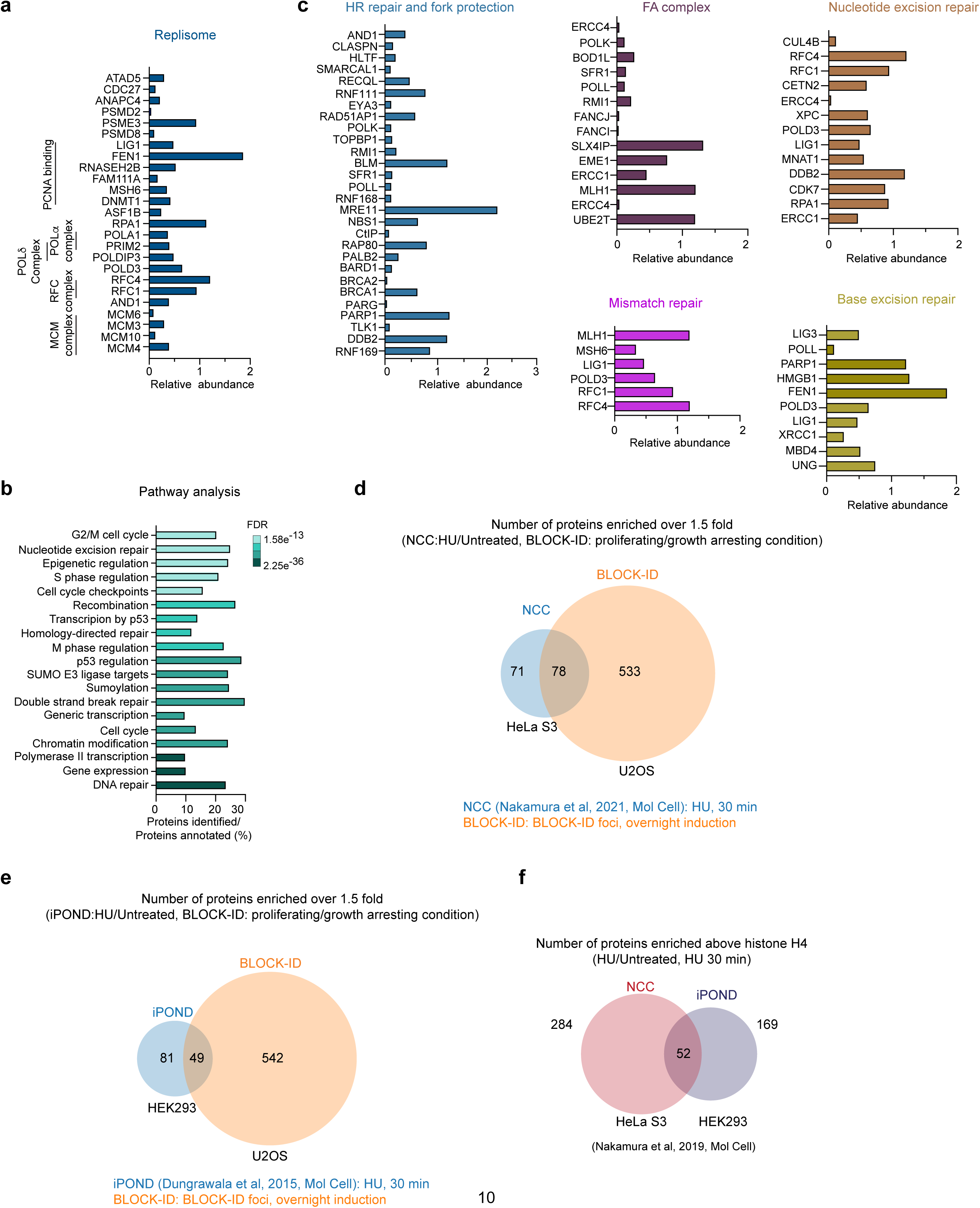
Replication and DNA repair factors identified by BLOCK-ID. **a,** Relative abundance of replisome-associated proteins identified by BLOCK-ID. **b,** Gene Ontology pathway analysis of proteins identified by BLOCK-ID. **c,** Plots showing the relative abundance of DDR factors identified by BLOCK-ID. **d,** Comparison of proteins identified by NCC^6^ and BLOCK-ID. **e,** Venn diagrams comparisons between proteins identified by iPOND^4^ and BLOCK-ID. **f,** Comparison of proteins identified by NCC and iPOND. Data in f were published in Nakamura et al 2019, *Mol Cell*^6^. Proteins in Venn diagrams of d and e are proteins with greater than 1.5 fold enrichment over controls.

**Extended Data Fig. 2:**
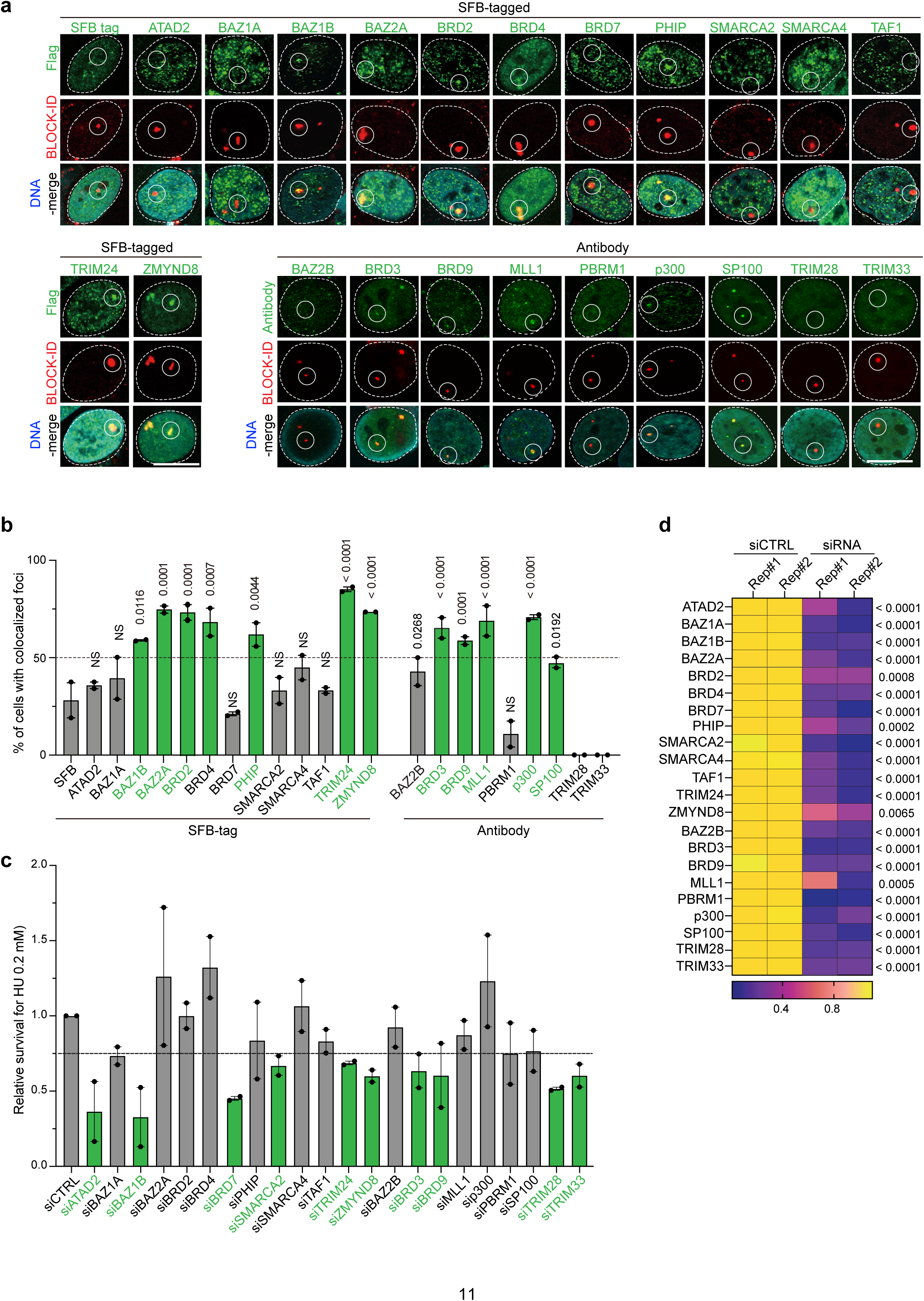
Validation of BRD proteins identified by BLOCK-ID. **a,** Screen for BRD protein localization to BLOCK-ID foci by IF and confocal microscopy. BRD proteins identified by BLOCK-ID MS were screened by either ectopic expression of SFB tagged BRD proteins or by antibodies. **b,** Quantification of BRD protein localization to BLOCK-ID foci. The horizontal dashed line represents the cut-off threshold for 50% of cells showing colocalization of BRD protein with BLOCK-ID foci, which are indicated in green. Statistical significance was determined with total > 30 cells per each experimental group in two biologically independent experiments. Statistical significance was determined using one-way ANOVA. **c,** Relative cell survival of U2OS cells treated with HU with the indicated siRNA treatment (see Methods). Cells were treated with the indicated siRNAs and cultured in 0.2 μM of HU for 14 days and surviving colonies were counted. Relative survival was normalized to non-targeting siRNA (siCTRL) cultured cells under the same condition. Horizontal dashed line represents 70% survival, with treatments leading to equal to or less than the cut off indicated in green. **d**, mRNA levels of siRNA treatments. RNAs isolated from U2OS cells post treatment of the indicated siRNAs were analyzed by qPCR and compared to siCTRL to determine the level of knockdown. Plot shows the extent of mRNA depletion in two biologically independent experiments. Statics significance was determined using one-way ANOVA. siRNAs were previously tested and validated in previous studies^38^. Unless otherwise indicated, error bars in graphs represent mean ± s.e.m., with P values indicated in the graphs. NS indicates no statistical significance. Dots on the graphs indicates independent experiments. N represents number of biologically independent experiments and indicated number represents total number of cells, or treats analyzed per each experimental group. All scale bars in IF images, 10 μm.

**Extended Data Fig. 3:**
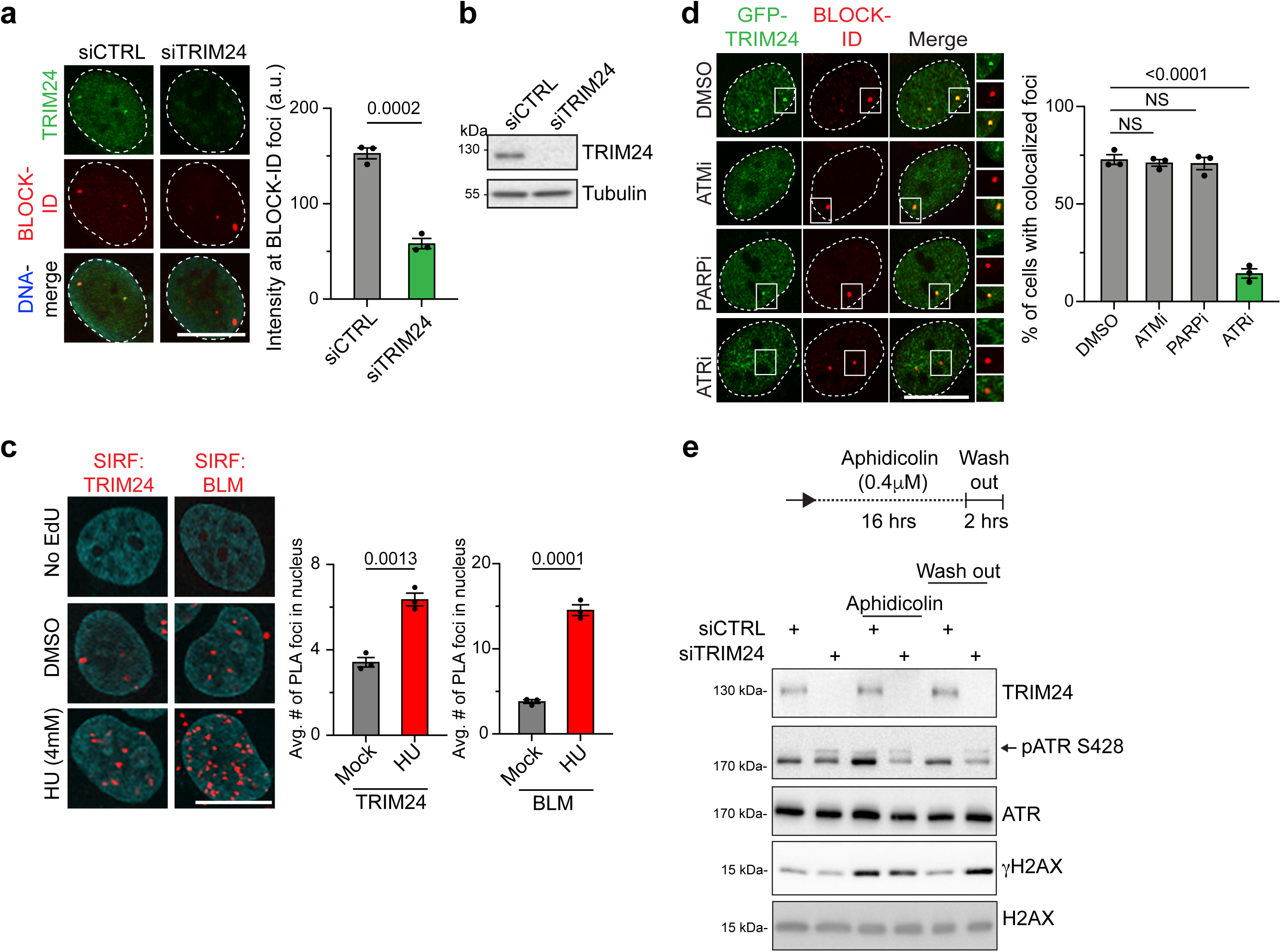
TRIM24 is involved in replication stress responses. **a,** Localization of endogenous TRIM24 at BLOCK-ID foci was analyzed by IF and confocal microscopy. Depletion of TRIM24 using siRNA (siTRIM24) reduced the intensity of TRIM24 antibody staining at BLOCK-ID foci and within the nucleus, validating the specificity of the antibody for IF. Quantification is on the right, n=3, >105. **b,** Western blot of lysates from **a** shows the depletion of TRIM24. **c,** SIRF analysis was performed to determine the proximity of endogenous TRIM24 or BLM to stalled replication forks following HU treatment. Cells were treated with DMSO or 4 mM HU for 3 hrs. No EdU and BLM act as negative and positive controls respectively for the assay. Quantification is on the right, n=3, >380. **d,** Recruitment of GFP tagged TRIM24 (GFP-TRIM24) to BLOCK-ID foci was analyzed by IF and confocal microscopy. Cells were treated with treated with 10 μM of KU55933 (ATMi), 1 μM of Olaparib (PARPi), or 1 μM of VE-821 (ATRi) for 14 hrs. Quantification with one-way ANOVA is on the right, n=3, >100. **e,** Design of experiment is illustrated on the top. Western blot analysis of lysates of U2OS cells from siCTRL and siTRIM24 treatment were probed with the indicated antibodies. Unless otherwise indicated, statistical analyses were performed with unpaired two-tailed T-test, error bars in graphs represent mean ± s.e.m., with P values indicated in the graphs. NS indicates no statistical significance. Dots on the graphs indicates independent experiments. N represents number of biologically independent experiments and indicated number represents total number of cells, or treats analyzed per each experimental group. All scale bars in IF images, 10 μm.

**Extended Data Fig. 4:**
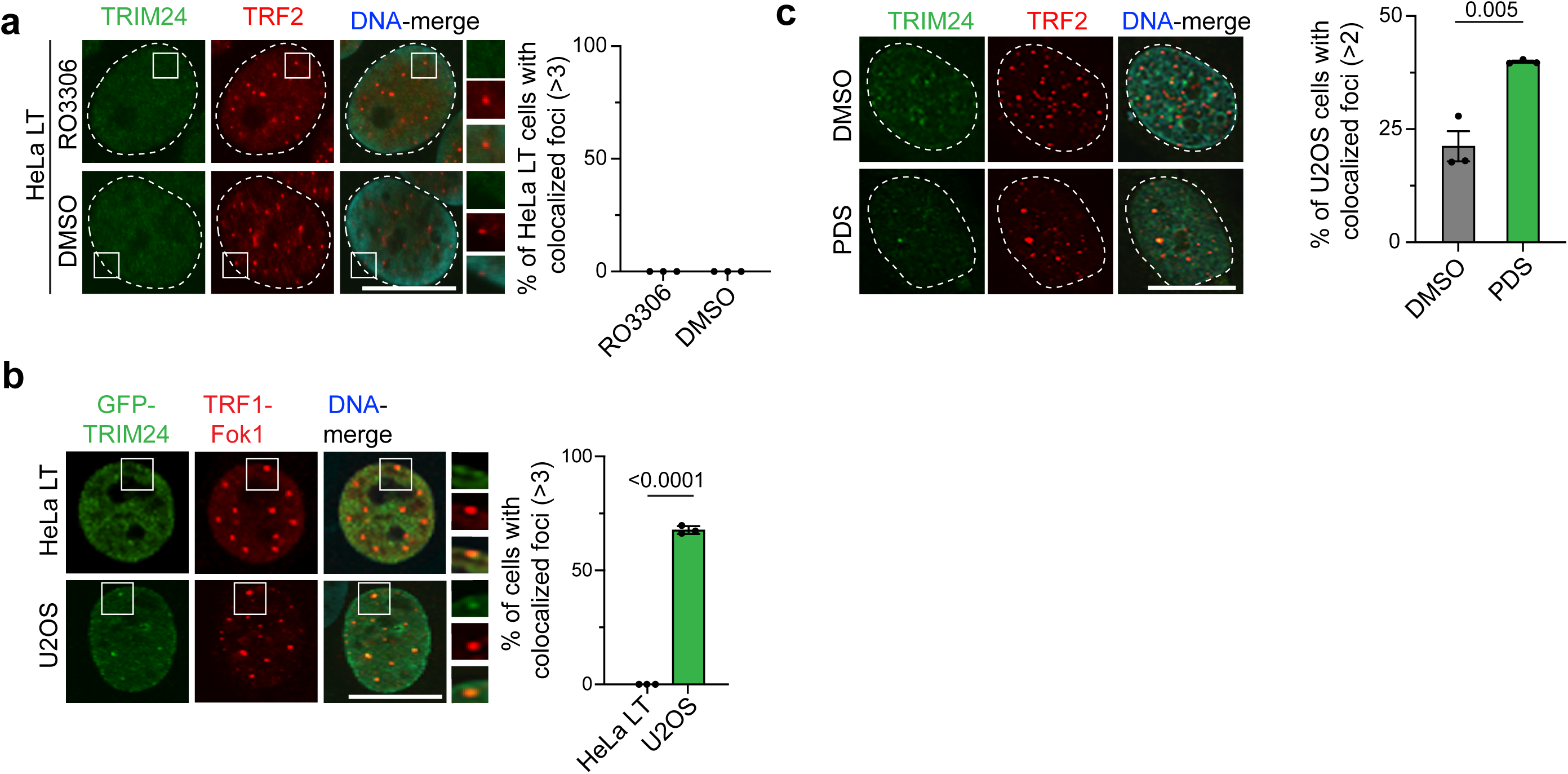
TRIM24 localizes to telomeres upon replication stress and in ALT. **a,** Localization of endogenous TRIM24 to telomeres was determined in HeLa LT cells by IF and confocal microscopy. Telomeres were detected using TRF2 antibodies. Cells were treated with DMSO or 9 μM of RO3306 for 16 hrs. Quantification is on the right, n=3, >110. **b,** Comparison of TRIM24 recruitment to TRF1-FokI induced DSBs in HeLa LT or U2OS cells was performed by IF and confocal microscopy. Quantification is on the right, n=3, >105. **c,** Localization of TRIM24 to telomeres in U2OS cells following treatment with pyridostatin (PDS) was determined by IF and confocal microscopy. Cells were treated with DMSO or 10 μM of PDS for 16 hrs. Quantification is on the right, n=3, >165. Unless otherwise indicated, statistical analyses were performed with unpaired two-tailed T-test, error bars in graphs represent mean ± s.e.m., with P values indicated in the graphs. NS indicates no statistical significance. Dots on the graphs indicates independent experiments. N represents number of biologically independent experiments and indicated number represents total number of cells, or treats analyzed per each experimental group. All scale bars in IF images, 10 μm.

**Extended Data Fig. 5:**
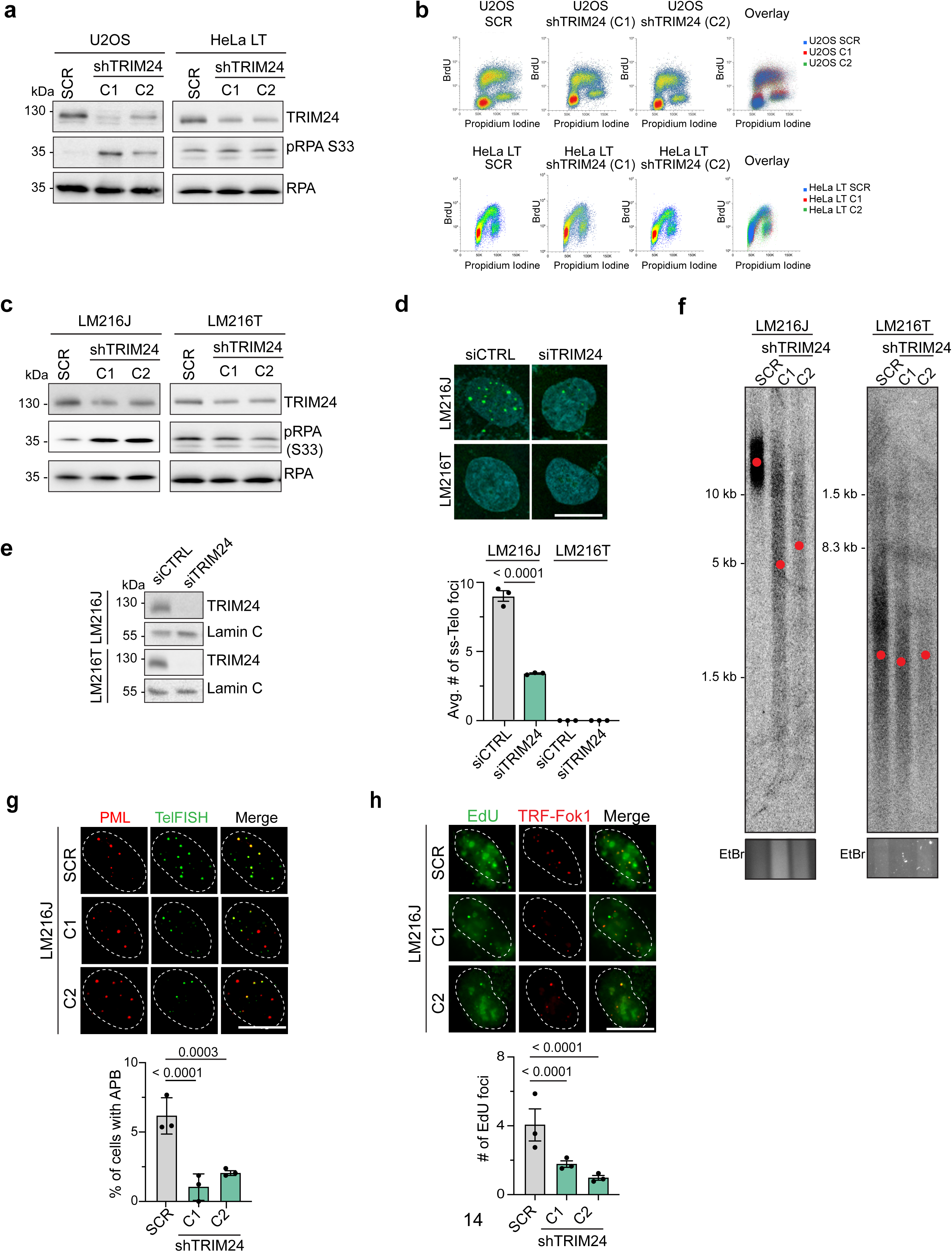
TRIM24 promotes telomere maintenance in ALT cells. **a,** Western blot analysis of shRNA treated U2OS and HeLa LT cells with the indicated antibodies. Cell lines were generated by stable expression of two independent shRNAs (C1 and C2) targeting the coding region of TRIM24. SCR is non-targeting scramble shRNA. **b,** Cell cycle analysis by flow cytometry of cells in **a** with two biologically independent experiments. **c,** Western blot analysis as in **a** in TRIM24-depleted LM216J (ALT+) and LM216T (ALT-) cell lines. Cell lines were generated by stable expression of two independent shRNAs (C1 and C2) targeting the coding region of TRIM24. SCR is non-targeting scramble shRNA. **d,** ssTelo foci were determined in LM216J or LM216T cells treated with siCTRL or siTRIM24 by FISH and confocal microscopy. Quantification is provided below, n=3, >830. **e**, Western blot of lysates from **d** using the indicated antibodies. **f,** Telomere length of the cell lines from **c** was determined using PFGE. Red dots indicate the mean telomere length. EtBr is a loading control. **g,** APB analysis was performed in TRIM24-depleted LM216J cells from **c**. Telomere FISH (TelFISH) labels telomeres. Quantification is provided below, n=3, >1500. **h,** Nascent telomere synthesis in TRIM24-depleted LM216J cells from c was analyzed. Telomere FISH (TelFISH) labels telomeres. Quantification is provided below, n=3, >90. Unless otherwise indicated, statistical analyses were performed with one-way ANOVA, error bars in graphs represent mean ± s.e.m., with P values indicated in the graphs. NS indicates no statistical significance. Dots on the graphs indicates independent experiments. N represents number of biologically independent experiments and indicated number represents total number of cells, or treats analyzed per each experimental group. All scale bars in IF images, 10 μm.

**Extended Data Fig. 6.**
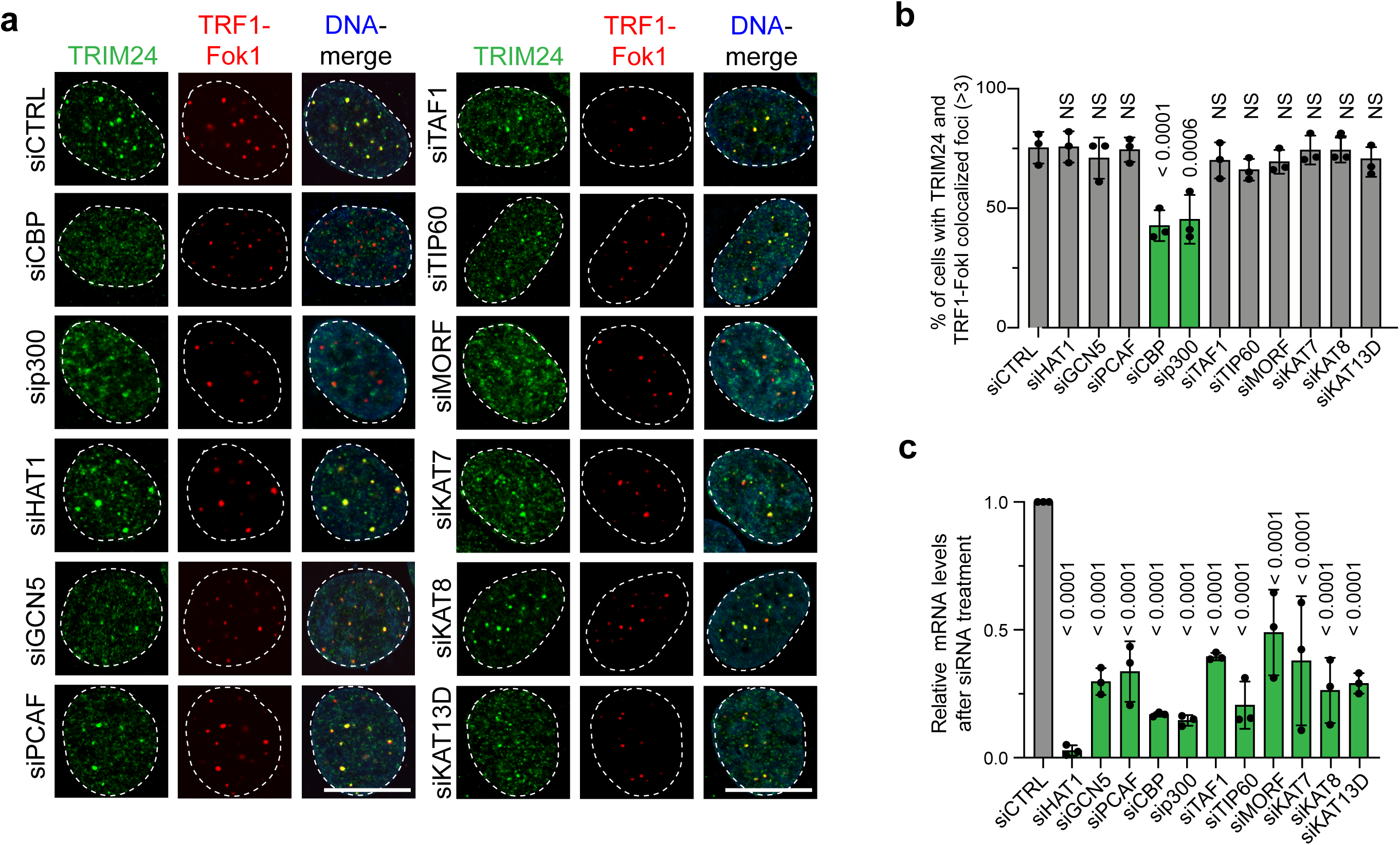
Screen for HATs required for TRIM24 localization to ALT telomeres identifies p300 and CBP. **a,** TRIM24 localization to TRF1-FokI labeled telomeres in U2OS cells was determined by IF and confocal microscopy following treatment with siRNAs towards human HAT genes as indicated. b, Quantification of a with > total 105 cells in three biologically independent experiments is shown. Statistic significance was determined using one-way ANOVA. **c,** Relative mRNA levels following treatment of the indicated siRNAs as analyzed by RT-qPCR. Statistic significance was determined using one-way ANOVA. Unless otherwise indicated, error bars in graphs represent mean ± s.e.m., with P values indicated in the graphs. Dots on the graphs indicates independent experiments. All scale bars in IF images, 10 μm.

**Extended Data Fig. 7:**
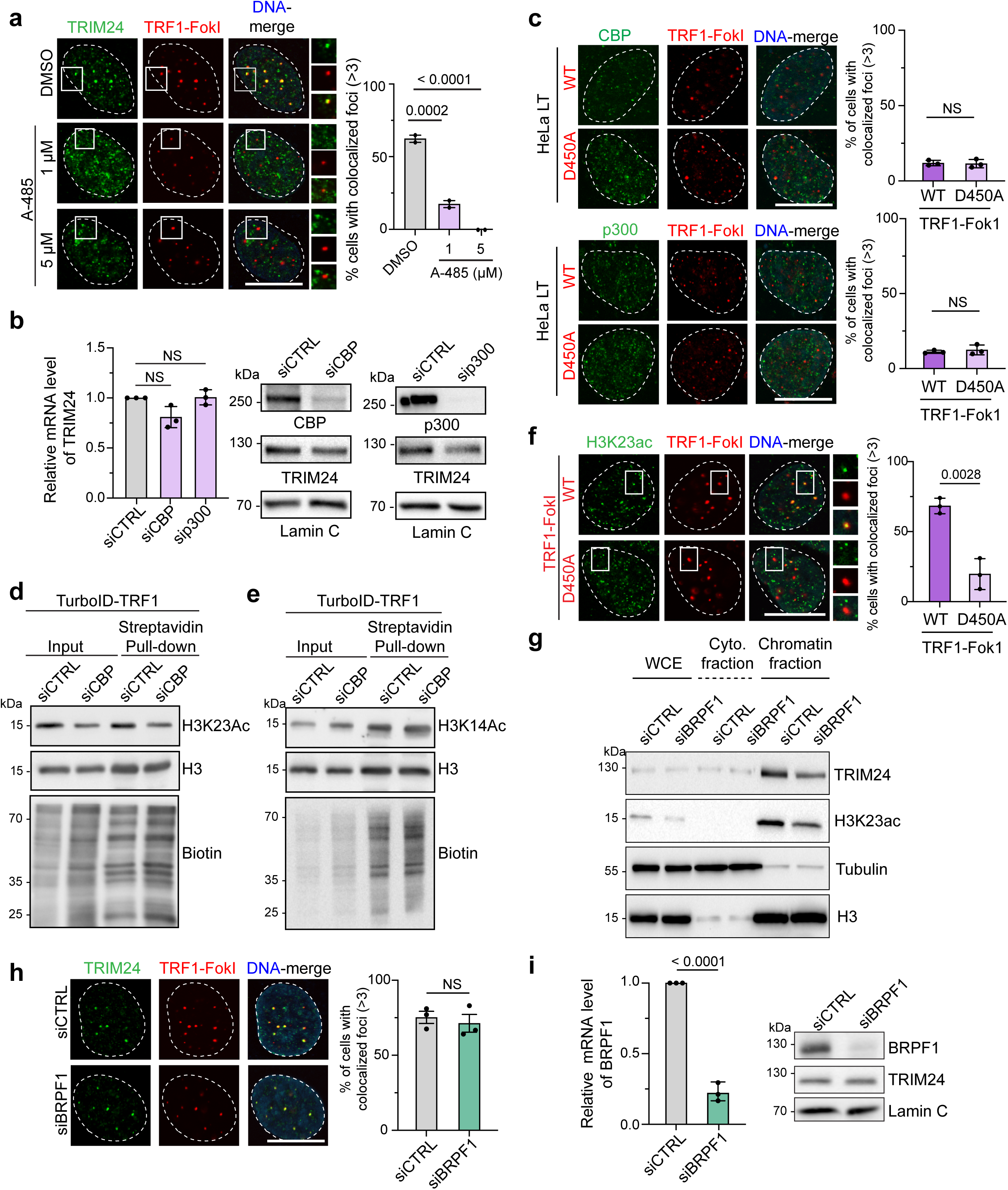
Regulation of TRIM24 telomere localization to ALT telomeres by p300 and CBP. **a,** Localization of TRIM24 to TRF1-FokI labeled telomeres in U2OS cells following treatment with the p300/CBP inhibitor A-485 was analyzed. Cells were treated with 1 μM or 5 μM of A-485 for 14 hrs. Quantification with one-way ANOVA is on the right, n=2, >123. **b,** mRNA levels of TRIM24 following CBP or p300 depletion by RT-qPCR. Statistical significance was determined using one-way ANOVA. Western blot of lysates of CBP or p300-depleted U2OS cells using siRNAs is shown on the right. **c,** Colocalization of endogenous CBP and p300 and TRF1-FokI WT or D450A labeled telomeres by IF and confocal microscopy in HeLa LT cells was analyzed by IF and confocal microscopy. Quantification of data is plotted, n=3, >104. **d-e,** Western blot of lysates from U2OS cells expressing TurboID-TRF1. The input and streptavidin purified proteins were analyzed by the indicated histone acetylation antibodies and biotin antibodies. histone H3 as loading control. **f,** Localization of H3K23ac at TRF1-FokI WT or D450A labeled telomeres in U2OS cells was analyzed by IF and confocal microscopy. Quantification is shown on the right, n=3, >60. **g,** Western blot analysis of chromatin fractionated lysates from U2OS cells following BRPF1 depletion. WCE: whole cell extract; Cyto: cytoplasmic fraction. **h,** TRIM24 telomere localization to TRF1-FokI labeled telomeres in U2OS cells following BRPF1 depletion was analyzed by IF and confocal microscopy. Quantification is on the right, n=3, >122. i, mRNA levels of BRPF1 by RT-qPCR following treatment with the indicated siRNAs. Western blot analysis of lysates from siCTRL and siBRPF1-treated U2OS cells with the indicated antibodies is shown on the right. Unless otherwise indicated, statistical analyses were performed with unpaired two-tailed T-test, error bars in graphs represent mean ± s.e.m., with P values indicated in the graphs. NS indicates no statistical significance. Dots on the graphs indicates independent experiments. N represents number of biologically independent experiments and indicated number represents total number of cells, or treats analyzed per each experimental group. All scale bars in IF images, 10 μm.

**Extended Data Fig. 8.**
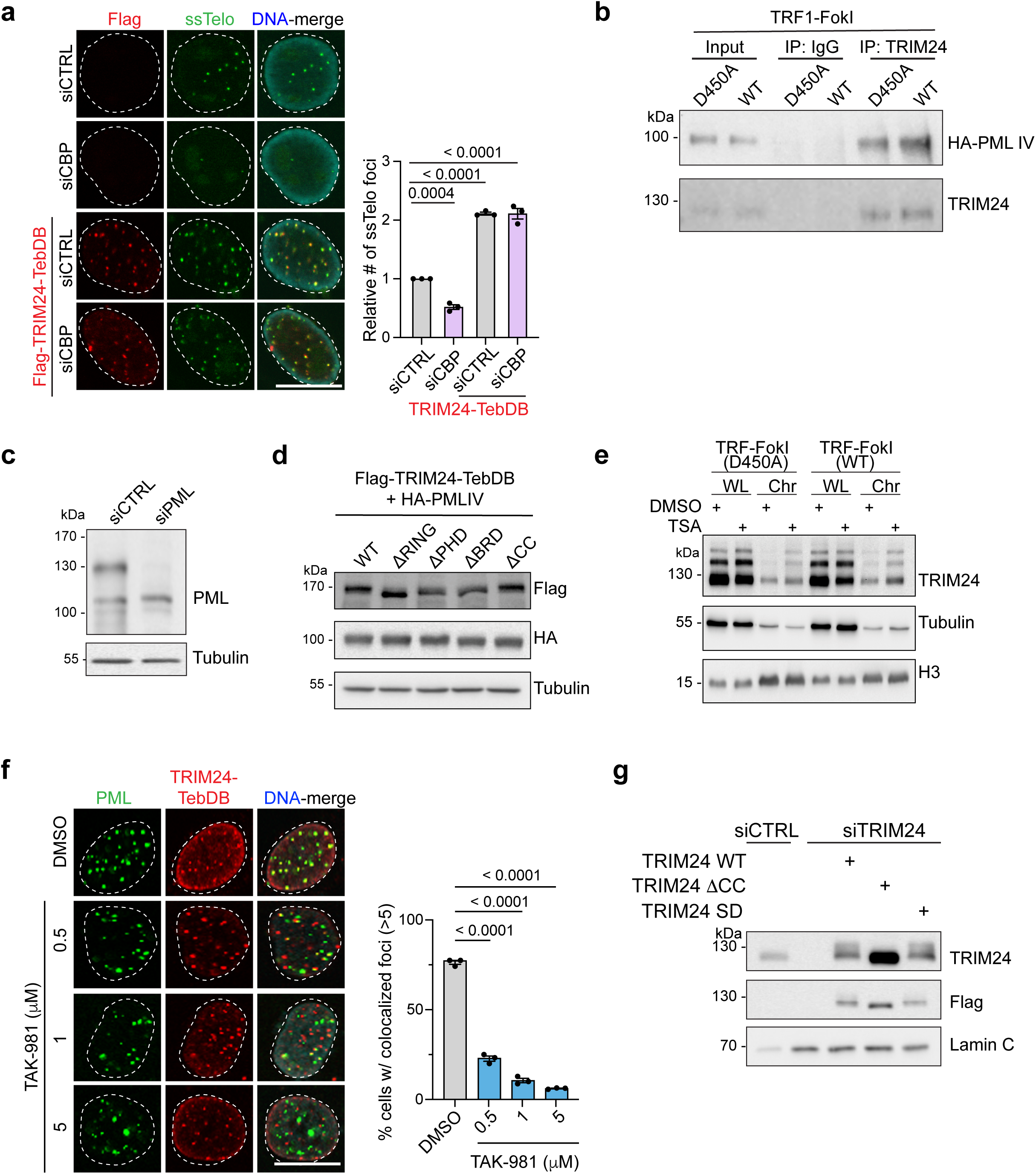
Identificaton of SUMO-dependent TRIM24 interaction with PML IV. **a,** ssTelo foci in U2OS cells were analyzed by by FISH, IF and confocal microscopy. Flag-TRIM24-TebDB was introduced to U2OS cells following CBP depletion using siRNAs. Quantification is on the right, n=3, >133. **b,** Interaction between endogenous TRIM24 and HA-PML IV in U2OS cells expressing TRF1-FokI WT or D450A was analyzed by immunoprecipitation. IgG serves as negative control. **c,** Western blot of lysates from siCTRL and siPML-treated U2OS cells determined knockdown of PML with Tubulin as loading control. **d,** Western blot analysis of lysates from U2OS cells expressing Flag-TRIM24-TebDB and its deletion variants and HA-PML IV. **e,** Detection of chromatin bound TRIM24 SUMOylation by Western blotting in U2OS cells expressing TRF1-FokI WT or D450A. Cells were treated with DMSO or 2 μM of TSA for 1 hr prior to sample isolation. WCE: whole cell extract, Chr: chromatin fraction. **f,** Colocalization of endogenous PML with Flag-TRIM24-TebDB was analyzed by IF and confocal microscopy in U2OS cells treatment with vehicle or TAK-981. U2OS cells were treated with the indicated concentration of TAK-981 for 16 hrs. Quantification is on the right, n=3, >93. g, Western blot of lysates from siCTRL and siTRIM24-terated U2OS cells complemented with TRIM24 and its variants. TRIM24 depletion was performed using a siRNA targeting the 3’ UTR of TRIM24. Unless otherwise indicated, statistical analyses were performed with one-way ANOVA, error bars in graphs represent mean ± s.e.m., with P values indicated in the graphs. NS indicates no statistical significance. Dots on the graphs indicates independent experiments. N represents number of biologically independent experiments and indicated number represents total number of cells, or treats analyzed per each experimental group. All scale bars in IF images, 10 μm.

**Extended Data Fig. 9.**
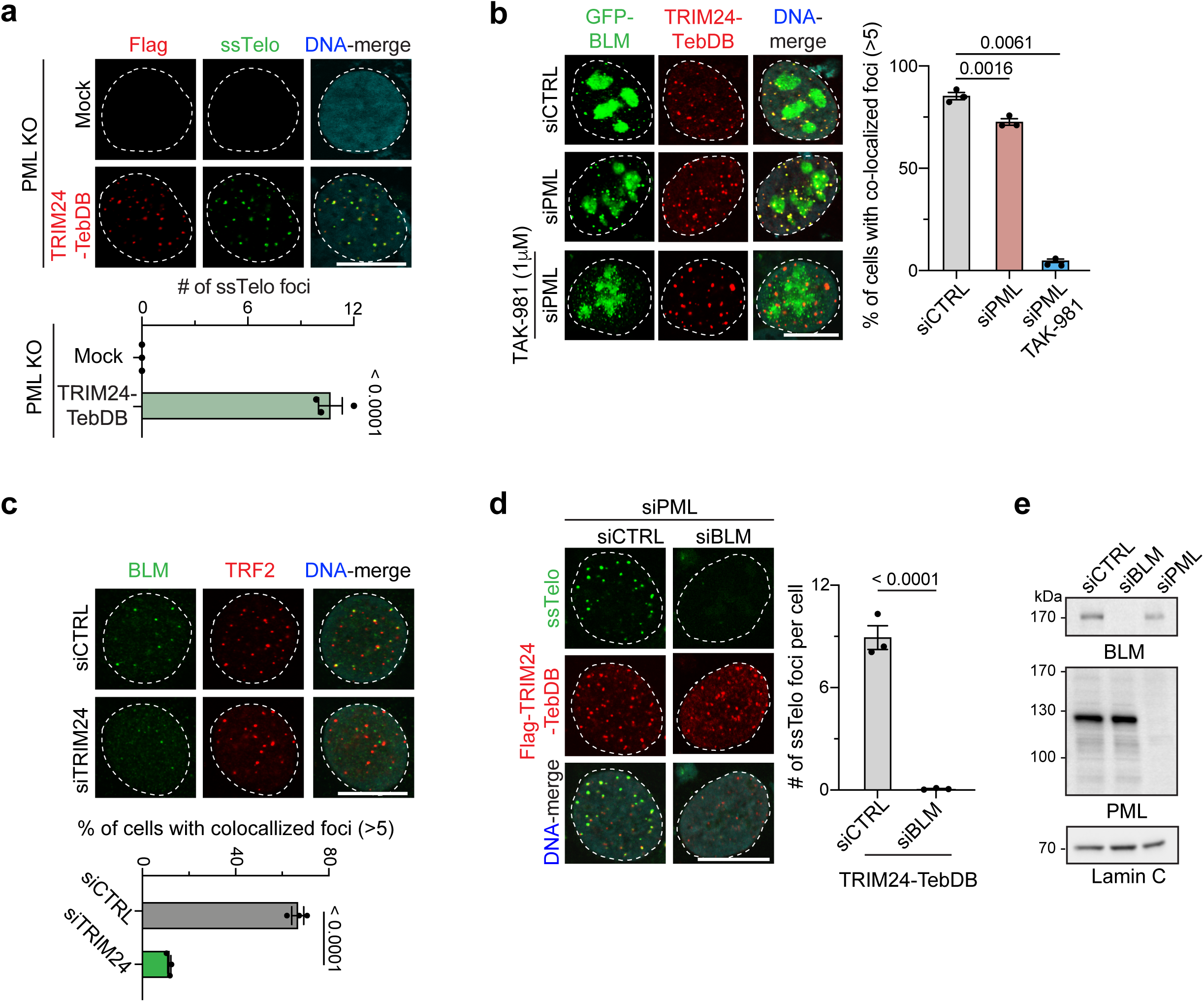
Analysis of telomere tethered TRIM24 in ALT. **a,** ssTelo foci in PML KO cells expressing Flag-TRIM24-TebDB was analzyed by FISH, IF and confocal microscopy. Quantification is on the right, n=3, >102. **b,** Recruitment of GFP tagged BLM (GFP-BLM) by Flag-TRIM24-TebDB in U2OS cells was analyzed by IF and confocal microscopy. U2OS cells were treated with siCTRL, and siPML. Following PML depletion using siRNAs, cells were treated with 1 μM of TAK-981 for 14 hrs. Quantification with one-way ANOVA is graphed on the right, n=3, >102. **c,** Localization of endogenous BLM to telomeres was analyzed by IF and confocal microscopy in U2OS cells treated with siCTRL and siTRIM24. Quantification is graphed below, n=3, >257. **d,** ssTelo foci were measured as in **a** in PML-depleted U2OS cells treated with siCTRL or siBLM that express TRIM24-TebDB. Quantification is on the right, n=3, >109. e, Western blot analysis of samples from **d** with the indicated antibodies. Unless otherwise indicated, statistical analyses were performed with unpaired two-tailed T-test, error bars in graphs represent mean ± s.e.m., with P values indicated in the graphs. NS indicates no statistical significance. Dots on the graphs indicates independent experiments. N represents number of biologically independent experiments and indicated number represents total number of cells, or treats analyzed per each experimental group. All scale bars in IF images, 10 μm.

**Extended Data Fig. 10.**
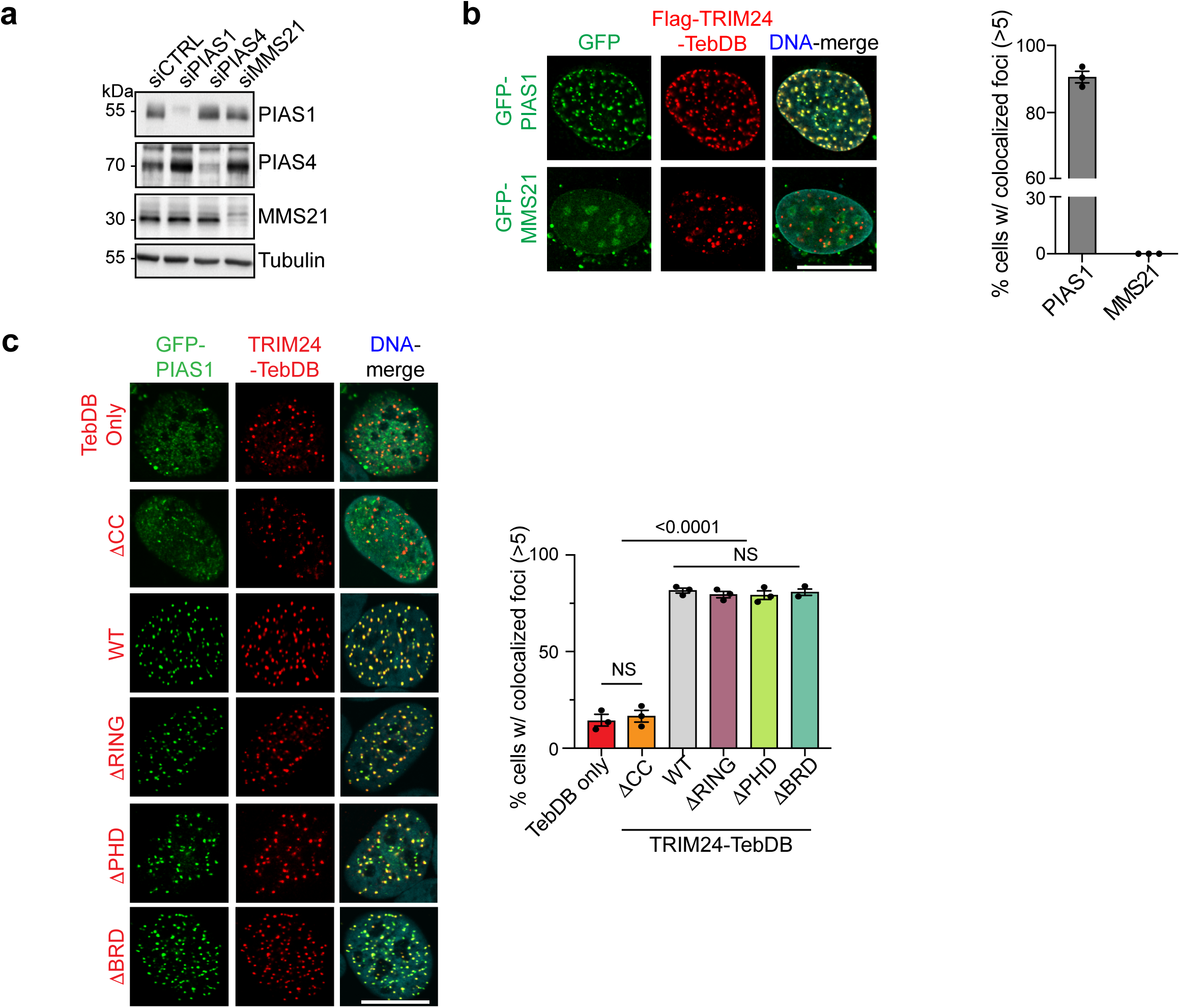
PIAS1 interacts with the cc domain of TRIM24. **a,** Western blot of lysates from U2OS cells treated with the indicated and probed with the antibodies listed. **b,** Recruitment of GFP tagged PIAS1 (GFP-PIAS1) or GFP tagged MMS21 (GFP-MMS21) by Flag-TRIM24-TebDB in U2OS cells was analyzed by IF and confocal microscopy. Quantification with unpaired two-tailed T-test is on the right, n=3, >93. **c,** Colocalization analysis between TRIM24-TebDB mutants and GFP-PIAS1 was performed using IF and confocal microscopy. Quantification is on the bottom, n=3, >96. Unless otherwise indicated, statistical analyses were performed with one-way ANOVA, error bars in graphs represent mean ± s.e.m., with P values indicated in the graphs. NS indicates no statistical significance. Dots on the graphs indicates independent experiments. N represents number of biologically independent experiments and indicated number represents total number of cells, or treats analyzed per each experimental group. All scale bars in IF images, 10 μm.

**Supplementary Table 1.** List of proteins identified by BLOCK-ID followed by mass-spectrometry analysis. Biotinylated proteins were selectively enriched using streptavidin biotin capture following induction of BLOCK-ID in proliferating and growth-arrested condition. Proteins exclusively identified in the proliferating condition, as well as those exhibiting a fold enrichment of greater than 1.5 times (based psm) are presented. The fold enrichment representing the ratio of protein abundance in proliferating versus growth-arrested conditions (normal/arrested) is expressed as Log2 value. “Unq.” indicates proteins uniquely identified in the proliferating condition.

**Supplementary Table 2.** List of siRNAs and shRNAs used in this study.

**Supplementary Table 3.** List of antibodies and their applications that are used in this study.

**Supplementary table 4.** List of qPCR primers used in this study.

